# Metabolic energetics underlying attractors in neural models

**DOI:** 10.1101/2023.03.11.532220

**Authors:** Richard B. Buxton, Eric C. Wong

## Abstract

Neural population modeling, including the role of neural attractors, is a promising tool for understanding many aspects of brain function. We propose a modeling framework to connect the abstract variables used in modeling to recent cellular level estimates of the bioenergetic costs of different aspects of neural activity, measured in ATP consumed per second per neuron. Based on recent work, an empirical reference for brain ATP use for the awake resting brain was estimated as ∼2×10^9^ ATP/s-neuron across several mammalian species. The energetics framework was applied to the Wilson-Cowan (WC) model of two interacting populations of neurons, one excitatory (*E*) and one inhibitory (*I*). Attractors were considered exhibiting steady-state behavior and limit cycle behavior, both of which end when the excitatory stimulus ends, and sustained activity that persists after the stimulus ends. The general finding was that the energy cost essentially follows the firing rate (average spikes/s) of the *E*-population, and firing rates of 8-10 spikes/s are in good agreement with the empirical reference value of ATP use. Self-sustained firing driven by recurrent excitation, though, involves higher firing rates and a higher energy cost. By considering a network in which each node is a WC model, we found that a combination of three nodes can serve as a circuit element that turns on when input passes a threshold and then persists after the input ends (an ‘on-switch’), with moderate overall ATP use. The proposed framework can serve as a guide for anchoring neural population models to plausible bioenergetics requirements.

## 1. Introduction

Ideas for understanding the complexity of brain function range from the cellular view, based on the cellular level complexity of individual neurons as information processing elements [1, 2], to the population view, based on how populations of neurons can function in ways that are qualitatively different from the individual neuron behavior [3, 4]. In the population view the biophysics of individual neurons is often highly simplified. The enormous success of artificial neural networks, modeling complex functions with very simple nodes (‘neurons’), suggests the potential power of complex network behavior to understand specific brain functions [5–7]. A recurring idea in neural population modeling is the potential role of neural attractors, an emergent feature of nonlinear dynamical systems [8].

Because the brain is a biological system, we also can look at brain function from the perspective of metabolic energetics [9]. Recent work at the cellular level has provided a quantitative picture of the energetic costs of multiple processes involved in neural function [10, 11]. The goal of the current paper is to begin to establish an energetics perspective in neural population modeling. Specifically, how do we use the results from cellular studies to attach energetic costs to the abstract and simplified mathematical components of population models? As an example, we consider the energetic costs of simple neural attractors based on the Wilson-Cowan model [12].

Biological systems are thermodynamic systems, and for physiological functions to happen in a reliable way they must be thermodynamically downhill (associated with an increase of entropy). For example, the distribution of sodium ions (Na^+^) across the cellular membrane, with a much higher concentration outside than inside the cell, is typically far from thermodynamic equilibrium. That is, for a negative membrane potential (more negative inside the cell than outside), the equilibrium distribution of sodium would be the reverse, with a higher concentration inside the cell. At an excitatory neuronal synapse, opening a sodium channel creates a strong inward positive current of Na^+^ ions, the basis of excitatory neural signaling, with a positive entropy change as the Na^+^ distribution moves closer to its equilibrium. Opening the sodium channel acts as a switch to enable a thermodynamically downhill pathway, a current of Na^+^ ions into the cell.

After the downhill process is completed, moving toward equilibrium, the system must be restored to its far from equilibrium state, a thermodynamically uphill process (decreasing entropy). For this to happen, the restoration processes must be coupled to another process with an even larger positive entropy change, so that the net process has a positive entropy change. In biological systems that strongly downhill process is primarily the conversion of adenosine triphosphate (ATP) to adenosine diphosphate (ADP). The ATP/ADP system is maintained far from equilibrium, with the concentration of ATP much higher than would be expected for equilibrium, providing a large entropy change when ATP is converted to ADP [13]. The ATP/ADP system must then be restored to its far from equilibrium state by reversing this process, converting ADP back to ATP. Again, this is thermodynamically uphill, and so must be coupled to another process with a stronger positive entropy change. In the brain, this process is the oxidative phosphorylation of glucose, creating ∼36 ATP coupled to the conversion of one glucose and 6 O_2_ molecules to water and carbon dioxide [14]. Blood flow delivers the glucose and oxygen to each part of the brain.

The ongoing energetic costs of neural activity can be quantified as the ATP consumed per second, which for a steady-state, can be related to the O _2_ consumed per second. The oxygen metabolic rate, in turn, relates to the signals measured with functional magnetic resonance imaging (fMRI), a widely used noninvasive tool for measuring activity dynamics in the human brain [15]. One motivation for this work is that a quantitative understanding of the energetic costs of model neural attractors will help to link the theoretical models with experimental fMRI measurements. In addition, models of the fMRI signal are central for estimation of neural responses and effective functional connectivity [16, 17], and these models could be improved by incorporating accurate estimates of the energetic costs. Our focus in this work is on estimating the energetic costs associated with neural population models.

## 2. Entropy and energy requirements underlying neural activity

### 2.1 Energy cost measured as ATP consumption

Neural signaling works, like many biological processes, by maintaining a system far from thermal equilibrium, with a ‘switch’ connected to the occurrence of a particular event that provides a pathway for the system to move toward thermodynamic equilibrium. For neural signaling, neurons are maintained far from equilibrium in several ways, including non-equilibrium ion distributions between inside and outside the cell, and neurotransmitter molecules packaged at high concentration in vesicles. Neural signaling is a thermodynamically downhill process, with arrival of an action potential as the initial switch. This initial event triggers a series of switches: calcium channels open on the pre-synaptic side of the synapse; calcium inflow triggers vesicle binding to the membrane and neurotransmitter release; and neurotransmitter binding to receptors on the post-synaptic neuron opens ion channels or engages second messenger systems and alters ion permeability. Recovery from this process, by restoring the different systems to their far from equilibrium state, requires coupling the restorative processes (e.g., pumping ions across the cell membrane, or recycling neurotransmitter and repackaging it in vesicles) to the dephosphorylation of ATP to ADP.

The energy cost of neural activity can be measured in terms of the rate of ATP use. However, it is good to keep in mind that by ‘energy cost’ we really mean ‘entropy cost’. That is, while the true energy changes associated with the conversion of one ATP to one ADP do not depend on the concentration ratio of ATP to ADP, the entropy generated by this conversion does depend strongly on that concentration ratio [18]. The strong positive entropy change comes from the fact that the ATP/ADP ratio is maintained at a level much higher than equilibrium. If this ratio decreases, there will be a point where the entropy increase associated with dephosphorylating one ATP molecule will be too small to offset the entropy reductions required for the restorative processes [9]. Because of the primary association with entropy, rather than energy, we can still think of the ‘energy costs’ in terms of the rate of use of ATP (ATP molecules/s converted to ADP molecules), but always with the assumption that the ATP/ADP concentration ratio is maintained at a normal level. In pathological states this assumption needs to be reconsidered.

### 2.2 The high energy cost of sodium ion currents

Estimates of the ATP required for different aspects of recovery from neural signaling suggest that the dominant cost is in pumping sodium out of the cell against its gradient [10, 19]. The sodium ion distribution is the system farthest from thermodynamic equilibrium, so the thermodynamic cost of restoring it is high (i.e., the equilibrium potential of the Na^+^ distribution is much farther from the resting potential than for the other ions). The sodium is moved across the membrane by the sodium/potassium pump, a molecule spanning the cell membrane, which couples the transport of 3 sodium ions out of the cell and 2 potassium ions into the cell with the consumption of one ATP molecule. This is thermodynamically uphill transport for both ions, indicating that the entropy increase available from one ATP/ADP conversion is much larger than the entropy increase needed to balance the transport of one of these ions.

One of the useful aspects of maintaining the sodium system so far from equilibrium is that it can act as a battery itself driving other systems. That is, sodium ion movement from outside to inside the cell is a thermodynamically downhill (entropy increasing) process, and it can be coupled to other uphill (entropy decreasing) processes such as pumping calcium ions against their thermodynamic gradient and recycling neurotransmitter. At least two transporters for calcium have evolved [20], one that couples entry of three sodium ions to the cell for each calcium ion transported out of the cell, and one that couples transport of one calcium ion to one conversion of ATP/ADP (ultimately the net energy cost is the same in terms of ATP use when pumping out the sodium is included). In short, the sodium gradient can act as a battery providing the entropy increase to drive other restorative processes, but ultimately the sodium gradient itself must be ‘recharged’ by coupling to ATP conversion with the sodium/potassium pump.

One way to think about the primary role of sodium currents in neural signaling is that opening a sodium channel acts as an amplifier of an arriving weaker signal in the form of neurotransmitter release [10]. This amplification is costly, in the sense that the energy cost of pumping back the sodium ions is much higher than the cost of recycling the neurotransmitter.

### 2.3 Experimental measurements of neural energy cost

To anchor the estimated energy costs in this paper to empirical metabolic rates we build on the recent work of Herculano-Houzel and colleagues [21, 22]. This work has re-shaped our view of the species differences of brain metabolism, illustrated with the data from [21] in **Figure 1**. The cerebral metabolic rate of glucose (CMRGlc) typically is measured as the metabolic rate of one gram or one cm^3^ of brain, and by this measure CMRGlc is more than 3x higher in mice compared to humans. Herculano-Houzel, using improved methods for cell counting, showed that the neuronal density varies substantially across species (indicated by the plotted point size in **Figure 1**). When this is taken into account, the glucose metabolic rate per neuron in the awake resting state is remarkably similar across several species of mammals. Note that this is the total metabolic rate of brain tissue per neuron, and so includes the costs of glial cells and other non-signaling cellular processes. In addition, not all metabolized glucose undergoes oxidative phosphorylation. For complete oxidation, the oxygen/glucose index (OGI, the ratio of O_2_ molecules consumed per glucose molecule consumed) would be 6. Hyder and colleagues [23], compiling data for a wide range of anesthetized states as well as the awake resting state found an average OGI of 5.6. Using that value, and assuming that full oxidation of a glucose molecule generates 36 ATP molecules, 1 μmole glucose/min would generate 3.3×10^17^ ATP/s. Dividing the glucose used by one gram of tissue by the number of neurons per gram gives the resulting ATP/s per neuron, plotted on the *y*-axis in **Figure 1**. As a reference, we define a typical energy use for the awake resting state as A_0_ = 2×10^9^ ATP/s-neuron. In the estimates to follow the ATP use of different aspects of neural activity are expressed as a multiple of A_0_.

**Figure 1.**
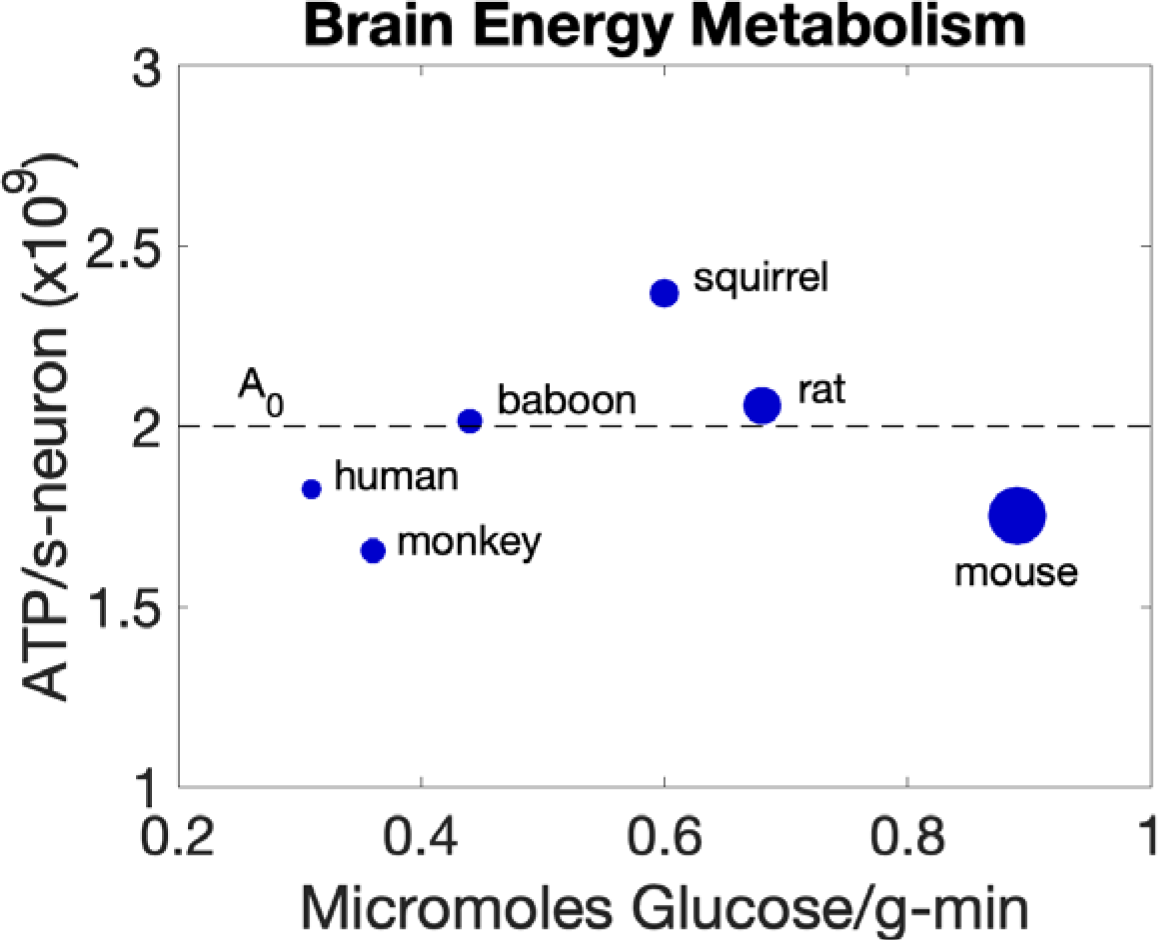
Whole brain average energy metabolism for different species. Based on glucose metabolism and neuronal density data from Herculano-Houzel [21]. On the *x*-axis is plotted the glucose metabolic rate per gram as typically measured. The diameter of each plotted point is proportional to the neuronal density for that species. From these data, the rate of ATP use per neuron was calculated and plotted on the *y*-axis. These calculations assumed an oxygen glucose index of 5.6 (i.e., an assumed fraction of 5.6/6.0 of glucose consumed was used in oxidative phosphorylation [23]), and that 36 molecules of ATP were produced from oxidative metabolism of one glucose molecule. The value A_0_ (dashed line) was taken as representative of awake resting energy metabolism for scaling the estimates in this paper.

A second useful anchor point is provided by the study of Hyder and colleagues [23]. In analyzing data for both the rat and humans under different levels of anesthesia they found that when neural signaling activity was reduced to near zero the glucose metabolic rate was reduced to about 25% of the awake resting rate. If we assume that this remaining energy cost represents the non-signaling cost needed to support neuron function, then this observation implies that in the awake resting state about 75% of the energy cost is due to neuronal signaling activity. More precisely, that 25% of baseline energy cost may represent the *minimum* absolute cost for non-signaling activity; that cost could increase in parallel with the signaling activity. In the following, we assume that 0.25A_0_ is the non-signaling energy cost, with the signaling energy costs calculated from the estimates for the neural models.

## 3. Estimates of energetic costs of neural activity at the cellular level

### 3.1 Assumptions

We are specifically interested in this paper in networks consisting of both excitatory (*E*) and inhibitory (*I*) neuronal populations. We also assume that action potentials (AP’s or spikes) generated by the *E*-population always arrive at excitatory synapses, and that AP’s generated by the *I*-population always arrive at inhibitory synapses. For a full energy cost estimate we assume that the non-signaling energy cost is 0.25A_0_, as described in the previous section.

Our basic hypotheses about the energetic costs of neural activity are:

1) The energy costs associated with generating and propagating an action potential (AP) are similar for the E and I populations, with a net energy cost proportional to the respective mean firing rates of the two populations.

In an influential paper, Attwell and Laughlin [10] made detailed estimates of the energy costs in terms of ATP use for different aspects of neural activity. They estimated the energy cost of an action potential by considering the charge needed to create a membrane potential corresponding to the peak of an action potential, given the assumed capacitance per unit area of the neuronal membrane and the total surface area of the cell and the axon carrying the AP. This required charge then defined how much Na^+^ is required to enter the cell to propagate the AP, and the ATP cost is then the cost of pumping this sodium back out. They estimated the minimum Na^+^ entry to be 2.9×10^8^ molecules. However, this estimate of the required charge is a minimum estimate for how much Na^+^ is needed. If a potassium channel opens, K^+^ will move out of the cell, reducing the accumulated charge inside. A critical component of ending an action potential is the opening of K^+^ channels. The central question for determining how much Na^+^ inflow is needed is then: how much overlap is there when both Na^+^ and K^+^ channels are open? Attwell and Laughlin, following earlier work on the squid giant axon, assumed that the amount of Na^+^ entry needed was 4x the minimum needed to charge the membrane due to charge loss through potassium channels. However, later work [24] showed that neurons are more efficient, with less overlap in time of Na^+^ and K^+^ fluxes. Howarth and colleagues [11] used these results to update the original estimates of the cost of forming and propagating an AP for cerebral cortex with a factor of only 1.24x the minimum Na^+^ needed. Based on that estimate, we assume that generating and propagating a single action potential requires entry of 3.6×10^8^ Na^+^. The energy cost of recovery is then 1.2×10^8^ ATP/spike to pump out the Na^+^. If the average firing rate of a neuron is one spike per second, then the energy cost in ATP/s-neuron would be 0.06 *A_0_*. Taking *F* as the average firing rate (in spikes/s) of a neuron in a population of neurons, the energy cost in ATP/s-neuron to generate and propagate action potentials is:

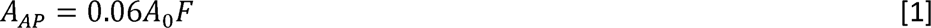

2) The pre-synaptic costs associated with arrival of an AP at a synapse (e.g., Ca^2+^ pumping, re-packaging neurotransmitter in vesicles, etc) are similar for both excitatory and inhibitory synapses.

We consider the presynaptic energy costs as those associated with the arrival of an AP at a synapse, including Ca^2+^ influx and vesicle release of neurotransmitter. In recovery, the Ca^2+^ must be pumped out, the neurotransmitter recovered through the action of the astrocytes, and the neurotransmitter re-packaged in vesicles. Attwell and Laughlin [10] made detailed estimates of these costs and estimated a net cost of 23,400 ATP are required in recovery from the pre-synaptic activity following the release of one vesicle of glutamate, with the assumption that a vesicle of glutamate contains 4000 glutamate molecules. We take the post-synaptic energy cost as the cost of pumping out Na^+^ and a smaller amount of Ca^2+^ that enter the post-synaptic neuron at an excitatory synapse after ion channels are opened. Attwell and Laughlin estimated this post-synaptic energy cost to be 140,000 ATP molecules per vesicle of glutamate released. Based on these estimates, the relative cost of presynaptic to post-synaptic activity is *a* = 0.17. These estimates of the pre-synaptic energy costs from Attwell and Laughlin were based on excitatory synapses, and we here assume that the same estimate applies for the pre-synaptic inhibitory synapses as well, with no post-synaptic energy costs for inhibitory synapses. Using the factor *a* the pre-synaptic energy costs can be anchored to the synaptic currents modeled in neural models (developed in the following sections).

3) Excitatory synaptic activity carries an additional post-synaptic energy cost required for pumping out the Na^+^ ions that entered through open Na^+^ channels in the course of excitatory signaling.

Attwell and Laughlin [10] estimated the energy cost of excitatory synaptic activity by estimating the number of synapses reached by each AP generated and considering the post-synaptic energy cost as discussed under Assumption 2. They estimated that a typical AP in the rat would release a total of 2,000 vesicles at the synapses with other neurons, for a recovery cost for the post-synaptic currents of 2.8×10^8^ ATP/spike. Here, though, we take a different approach that is tied directly to the way synaptic currents are treated in neural models (developed in detail in the next section). The essential difference in the approach here is that Attwell and Laughlin assumed a value for the average firing rate that was not directly tied to the excitatory synaptic currents. Here we want to close that loop by considering that the average net synaptic current per neuron in a population determines the average firing rate of the neurons in the population. This will allow us to interpret the more abstract representation of synaptic currents in neural models in terms of physiological currents and associated energy costs. The factor *a*—the ratio of pre-synaptic energy costs to post-synaptic energy costs—then provides an estimate of the pre-synaptic energy costs associated with a given synaptic current.

### 3.2 Synaptic current needed to initiate an action potential

#### Neural spiking

A basic simplified model of how a neuron works, often used in neural modeling, is as follows [12]. Synaptic currents, if strong enough, lead to a neuron firing an action potential (a spike). For a time *τ* after this the neuron cannot fire again (i.e., the maximum spiking rate is 1/ *τ*). If*S* is the net synaptic current, the firing rate *F* of the neuron (spikes/s) is modeled as:

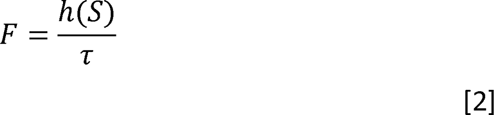

where *h(S)* is a nonlinear sigmoidal response function, with *h(S)*=0 if*S*<0 and *h(S)* approaches 1 as *S* gets very large. Between these two extremes *h(S)* increases monotonically, and we define *S_0_* as the value for which *h(S_0_)*=0.5. Here *F* is not necessarily a periodic firing rate, but rather the probability per unit time that the neuron will fire (i.e., the probability of firing in any short time interval *dt* is *Fdt*). There are several components of Eq [2] that we need to interpret in specific physiological terms: the synaptic currents (*S*), the response function *h(S)*, and the time constant *τ*.

#### Synaptic currents

Our basic model of neural spiking is that when the membrane potential at the soma increases by a threshold value *V_th_* an action potential (AP) will fire [25]. The change in potential depends on the net input current. This current depends on an excitatory inward positive current due to entering Na^+^ ions at excitatory synapses (*I_E_*), and on an inhibitory current opposing the positive Na^+^ current due to inhibitory synaptic activity (opening K^+^ and Cl^−^ channels). We treat the latter as equivalent to an opposing current *I_I_*, so that the net inward positive current is *I* = *I_E_* – *I_I_*. Note that the direct effect of synaptic activity is to alter ion conductance by opening specific channels. The increased conductance leads to currents that are proportional to the product of the conductance and ( *V* - *V_R_*), the difference in membrane potential between the current value *V* and the reversal membrane potential for that ion ( *V_R_*, the potential for which the distribution of that ion would be at equilibrium). Because the reversal potential of Na^+^ is much more positive than the resting potential, increasing Na^+^ conductance will quickly lead to an inward positive current. Inhibitory synaptic activity could create an outward positive current for K^+^ channels opening, because the K^+^ reversal potential is less than the resting potential. For Cl^−^ channels the reversal potential is near the resting potential, so initially there may be no associated current. However, as the membrane potential depolarizes (becomes more positive), an inward Cl^−^ current opposing the positive Na^+^ current will develop.

#### The effective refractory period *τ*

Following other modeling papers [12, 25], our interpretation of *τ* is that it can be longer than the classical absolute refractory period associated with an action potential (∼2 ms), and that the time *τ* should be viewed as an effective refractory period for the population behavior.

#### The firing rate

Our basic model of neural spiking, following the modeling in [25], is that once the AP is generated the net charge that has built up in the cell is dissipated. Then the neuron begins to recharge at a rate that depends on the net input current *I*. If *T_th_* is the time required for the membrane potential to change by the threshold value *V_th_*, the time (on average) between AP’s is *τ* + *T_th_*. The probability per unit time of generating an AP (the average firing rate *F*) is:

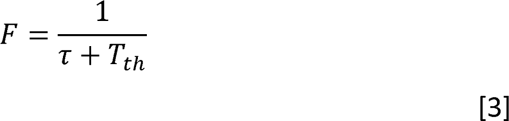

If *T_th_*<<*τ*, the firing rate approaches its maximum value 1/ *τ*. In **Appendix 1**, a simple model for *T_th_*is described based on a leaky integrate and fire model. The parameters of the model are: the voltage threshold *V_th_* required to initiate an AP; the input membrane resistance *R*; and the membrane capacitance *C*. The critical parameter for determining the firing rate is the threshold current *I_th_=V_th_/R*. To anchor the modeling to experimental data, we used the results of Degenetais and colleagues [26], who measured several cellular properties in prefrontal cortex pyramidal neurons in the rat. They tested for the lowest current applied in a 400 ms pulse that would evoke an AP, finding an average of 0.3 nA. Interestingly, they also measured the voltage threshold (17 mV) and the input resistance (34 MOhm). For these measurements, from the model expression (*V_th_/R*) the threshold current is about 0.5 nA. This difference (between 0.3 and 0.5 nA) may reflect the challenges of making accurate measurements of *R*, or the limitations of the leaky integrate and fire model applied to actual neurons. For our energy cost estimates here we assume *I_th_* =0.3 nA based directly on the measured result (firing an AP) rather than the inferred value from the model. Nevertheless, the model is a useful way to connect firing rates to input current anchored by that empirical value for *I_th_* (as shown in **Figure 2**). If the input current is less than *I_th_*, the threshold potential for spike generation will never be reached and the firing rate is zero. Above the threshold the firing rate initially rises very rapidly, as illustrated in **Figure 2**.

**Figure 2.**
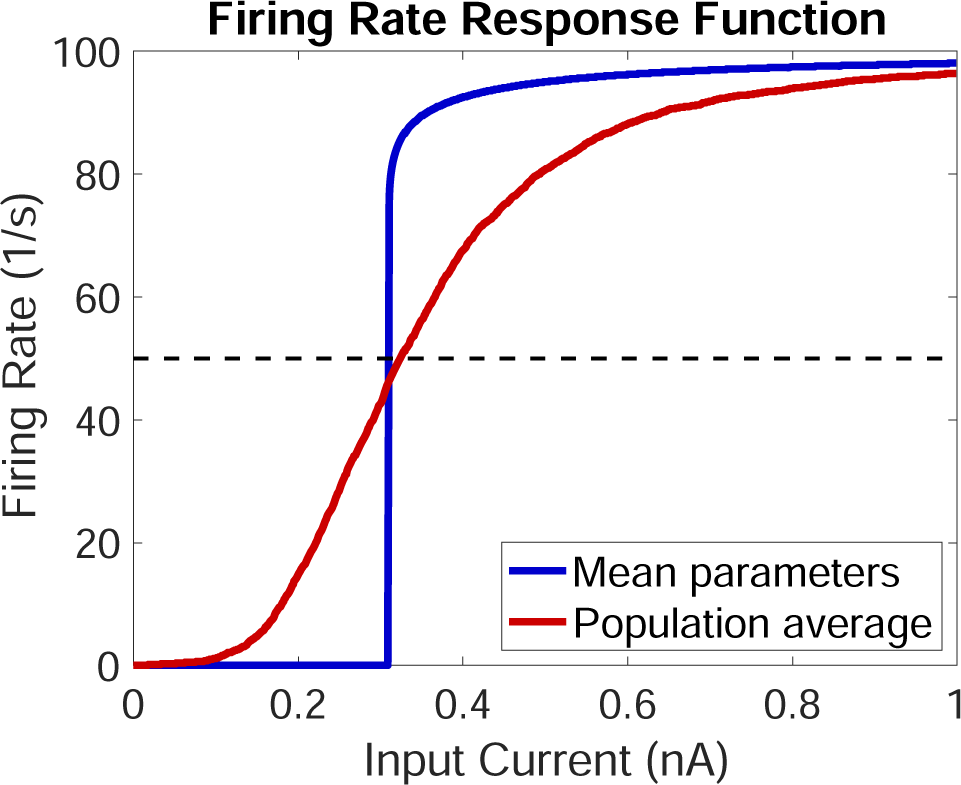
Model for the firing rate response function. Firing rate as a function of input current (blue) calculated with the model described in the **Appendix** with a threshold current for initiating an action potential *I_th_* = 0.3 nA. The average population response (red) when each of the cellular parameters was drawn from a Gaussian distribution with standard deviation 30% of the mean, creating a sigmoidal average response function.

#### The sigmoidal response function *h(S)*

In the neural modeling considered here we are specifically interested in average population behavior, rather than individual neuron behavior, and so need to consider variations between the neurons. To estimate the average response for a variable population of neurons, the cellular parameters *V_th_*, *R* and *C* were each assumed to vary independently across neurons with a Gaussian distribution with standard deviation 30% of the mean. For each parameter a value was chosen from their respective distributions and the corresponding response curve was calculated. Curves were calculated for 3000 such choices and the response curves averaged to give the resulting sigmoidal shaped average response curve (the red curve in **Figure 2**).

#### Connecting synaptic activity in neural models to currents

In the neural models we are considering, the firing rate response function is assumed to be of the form of Eq [2]. The synaptic current is represented by a variable *S*, and the mathematical form chosen for *h(S)* is usually a relatively simple smooth curve with a sigmoidal shape (such as the red curve in **Figure 2**). The key for anchoring the mathematical model to physical currents is to identify the current corresponding to *S_0_*, the value that produces a firing rate half of the maximum. Based on the simulations in **Appendix 1**, and illustrated in **Figure 2**, we assume that the model synaptic current *S_0_* approximately corresponds to the threshold current *I_th_* (in the simulations of **Figure 2**, the current corresponding to half-maximum firing is actually slightly larger, 1.08 *I_th_*, and depends slightly on the membrane capacitance; given the large uncertainties in estimating *I_th_*, we neglect this slight difference). Our basic assumption is then that the physical current equivalent of *S=S_0_* is the assumed threshold current *I_th_*= 0.3 nA.

#### Energy cost of synaptic currents

At this point it is useful to consider the net synaptic current *S* as the difference of an excitatory current *S_E_* due to Na^+^ entering at excitatory synapses, and an opposing inhibitory current *S_I_*due to the inhibitory synapses: *S* = *S_E_* – *S_I_*. The firing rate is determined by the net synaptic current *S*, but by our assumption 3 the energy costs in recovery from those currents are due primarily to the excitatory current *S_E_*. To connect a Na^+^ current to the model parameter *S*, consider first the case when there is no inhibitory synaptic current: how much Na^+^ current is needed to produce a synaptic current equivalent to *S_0_*? One coulomb of charge is equivalent to 6.24×10^18^ Na^+^ ions, so the current corresponding to *S_E_* = *S_0_* is 1.87×10^9^ Na^+^/sec. Recovery, clearing the accumulated Na^+^, requires 6.2×10^8^ ATP/s, or 0.31A_0_. The energy cost of excitatory synaptic currents (in ATP/s-neuron) as represented in the modeling framework is:

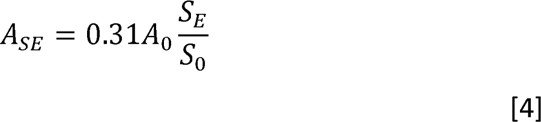

Finally, we return to the earlier estimate of the pre-synaptic energy cost, which we assume applies to both excitatory and inhibitory synapses. Based on the estimate from Attwell and Laughlin [10], at an excitatory synapse the pre-synaptic cost is 17% of the post-synaptic energy cost. The pre-synaptic energy costs for excitatory currents *S_E_*and inhibitory currents *S_I_* are then:

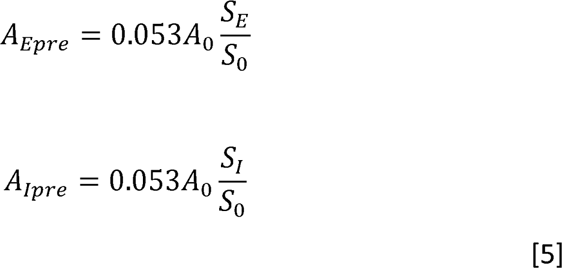

In summary, for a sigmoidal response function *h(S)*, with *S_0_*the half-maximum value of *h(S)*, the firing rate is given by Eq [2], and the associated energy costs are given by Eqs [1, 4 and 5].

## 4. The Wilson/Cowan model of neural population dynamics

The Wilson-Cowan (WC) model has been used in many theoretical studies since it was first introduced in 1972 [12, 27–34], often because it is an example of how oscillations can arise just from nonlinear population dynamics even though the underlying neurons are not intrinsically oscillatory. The WC model describes the bulk behavior of a population of excitatory neurons (‘*E*-neurons’) interacting with a population of inhibitory neurons (‘ *I*-neurons’), illustrated in **Figure 3A**. Here our goal is to use the model to generate examples of nonlinear behavior and investigate the associated energy costs.

**Figure 3.**
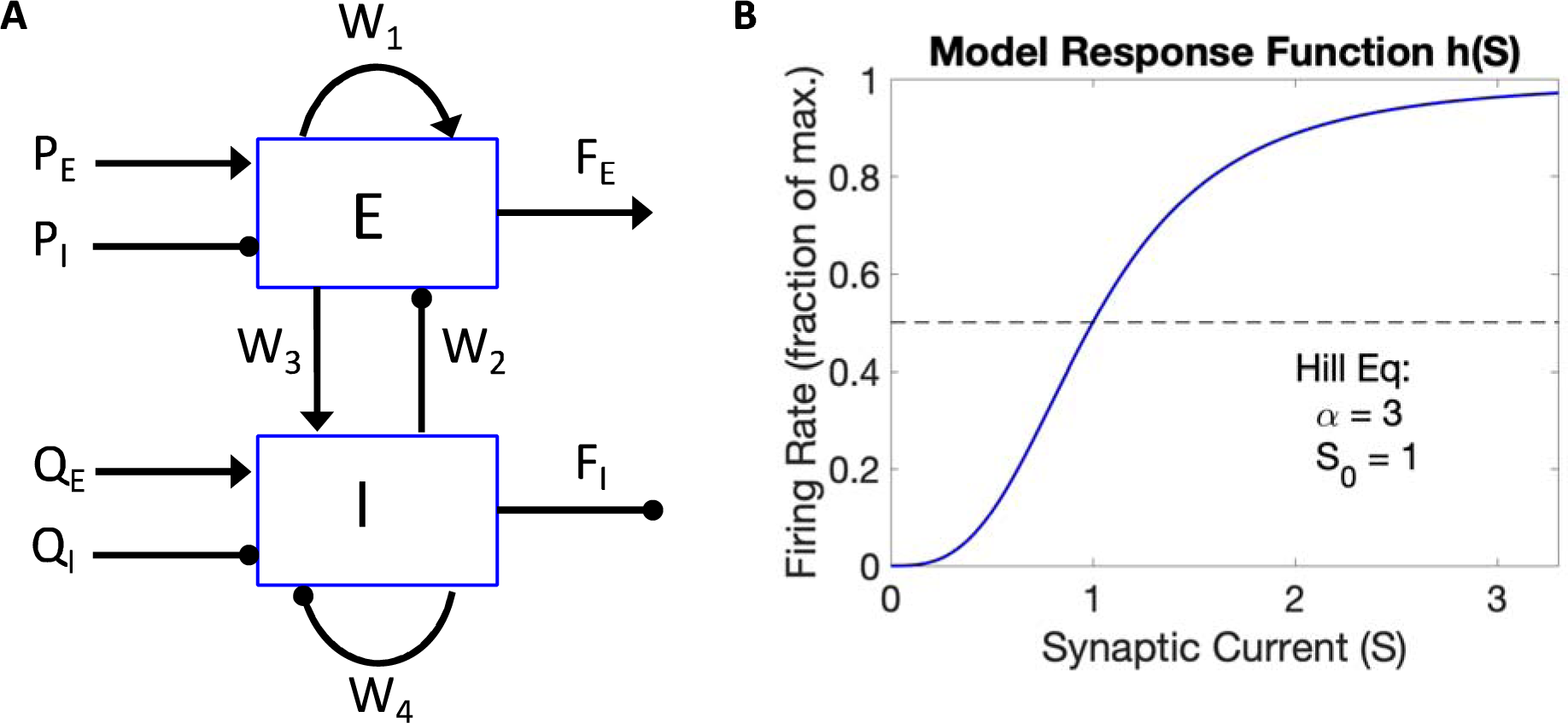
Wilson-Cowan model. **A**) A model of two interacting populations of neurons, one with excitatory outputs ( *E*) and one with inhibitory (*I*) outputs. Connectors ending with an arrow represent excitatory inputs, and those ending with a disk represent inhibitory inputs. The internal weights *W* describe the population interactions, and the four *P* and *Q* variables are external inputs. The outputs of the model are the average firing rates of the two populations ( *F_E_* and *F_I_*). **B)** The assumed firing rate response function *h(S)* as a function of net synaptic input *S*, based on the Hill equation (Eq [6]).

We focus on three types of neural attractor, which we describe as: steady-state responses, limit cycle behavior, and persistent self-sustained activity. All three have the property of an attractor that, following a brief perturbation, the system settles back to the attractor state. The two attractors that we refer to as a steady-state response and persistent activity could be described as point attractors, differing in how they depend on the excitatory input to the model. Steady-state behavior depends on the magnitude of the input, and ends when the input ends. Limit cycle behavior occurs when the steady-state becomes unstable, leading to cyclic oscillations of activity until the input ends. Persistent activity occurs when the recurrent excitation within the population leads to a self-sustained constant high firing rate after the input has ended. The latter depends on a threshold magnitude of the input current, initiating when the threshold has been exceeded and then persisting at an activity level related to the population dynamics rather than the input magnitude. We then consider neural networks in which each node is a WC-unit, and describe an energy efficient neural circuit element (composed of a few nodes) that serves as a ‘neural on-switch’, turning on an output node, with sustained activity at a moderate firing rate, when the input to the input node exceeds a threshold value.

### Assumptions

For the mathematical form of the basic equations we essentially follow Wallace and colleagues [27]. We consider a population of *N_E_* excitatory neurons interacting with a population of *N_I_* inhibitory neurons, and assume that 85% of the neurons are excitatory (*N_E_*/(*N_E_*+*N_I_*)=0.85). For each population there is an effective refractory period *τ* after generating an AP when the neuron cannot fire again. Here we assume *τ*_E_ = 10 ms and *τ*_I_ = 7 ms, similar to previous work [12]. For each population the average firing rates follow Eq [2]. The sigmoidal response function *h(S)* is assumed to be the same for each population, and we used the Hill equation:

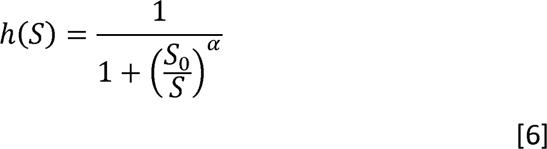

with the half-maximum value *S_0_*=1 and Hill exponent *α*=3 (illustrated in **Figure 3B**).

### Synaptic currents

For the two interacting populations of *E*- and *I*-neurons, we define the variable *E(t)* as the fraction of the *E*-neurons that are in the refractory state at time *t*, and a similar definition of *I(t)* for the *I*-neurons. Physically, *E(t)* and *I(t)* are then proportional to the fraction of the respective populations that fired during an interval *τ* in the recent past, and so these activity levels provide measures of the number of arriving spikes producing synaptic inputs from the two populations. We define four synaptic currents: *S_EE_* is the excitatory synaptic current per neuron in the *E*-population; *S_EI_* is the inhibitory synaptic current for *E*-population neurons; *S_IE_* is the excitatory synaptic current for *I*-population neurons; and *S_II_*is the inhibitory current to the *I*-population neurons. In this notation the first subscript indicates the population, and the second indicates whether it is excitatory or inhibitory current. These synaptic currents are modeled as:

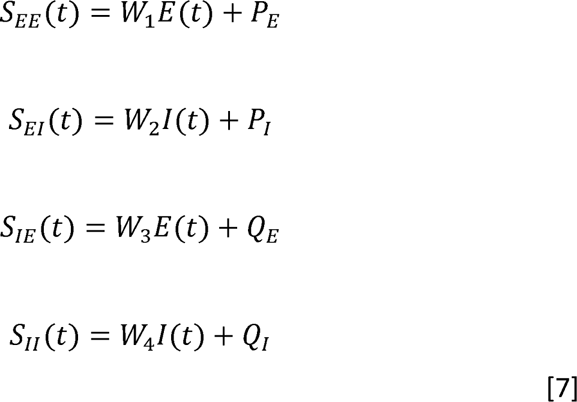

Each of the two populations connects to itself and to the other population through the weights (*W*’s) (diagramed in **Figure 3A**). Each population also has an external excitatory input (*P_E_* and *Q_E_*) and an external inhibitory input (*P_I_* and *Q_I_*). (The *P* and *Q* notation goes back to the original paper [12], as an input that could be either positive or negative; here they are separated into excitatory and inhibitory because excitatory input entails a higher energy cost.) Note that the synaptic weights *W* are model parameters that we do not have to define explicitly in physiological terms in order to estimate energy costs. That is, the energy costs are assumed to just depend on the magnitude of the synaptic currents produced (i.e., the connection of *S=S_0_* in the model to a corresponding physical current). A weight *W* could be increased in several ways: adding more synapses, increasing vesicle release probability, increasing the channel open time, etc. For modeling learning, the weights are subject to modification related to ongoing activity.

### Population dynamics

The average firing rates of the two populations at time *t* are:

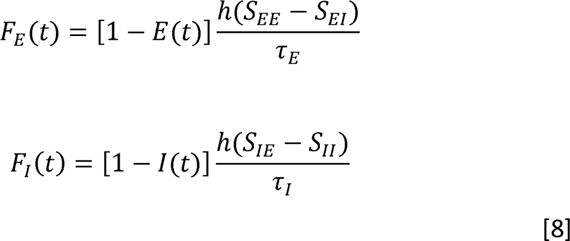

Neurons are either in the refractory state, having recently fired, or in a quiescent state ready to fire. The first term in brackets in each expression is the fraction of the population able to generate an AP. As the two populations interact under the influence of the external drivers, the rate of change of *E* and *I* are modeled as the difference between the rate at which neurons in the refractory state are emerging from that state ( *E*/ *τ*) and the rate at which the quiescent neurons are entering the refractory state by firing:

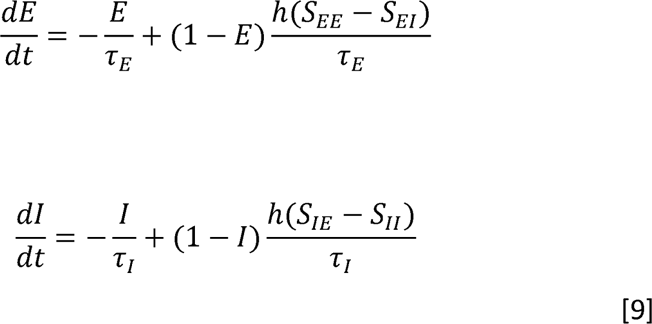

### Calculation of model dynamics

A particular model is specified by the values of the four weights *W*, the two time constants *τ*_E_and *τ*_I_, and the parameters of the firing rate function *h(S)*. For given dynamic inputs to the model (*P_E_*, *P_I_*, *Q_E_*, and *Q_I_*), Eq [9] is solved numerically for*E(t)* and *I(t)*. The corresponding firing rates of the two populations, *F_E_(t)* and *F_I_(t)* are then calculated from Eq [8]. In this paper we focus on reporting those firing rates under different conditions. It is useful to note that these values are the average values for the population. Although the maximum firing rate for an individual neuron in the *E*-population is 1/ *τ*_E_, the maximum steady-state average for the population is half that due to the factor of (1-*E*) in Eq [8].

### Energy costs of the model network

The energy costs are estimated based on Eqs [1, 4, and 5]. The 3 categories of spiking, pre-synaptic activity, and post-synaptic activity are combined separately for the E- and I-populations as *A_E_* and *A_I_*, in each case expressed as ATP/s-neuron of that population. The reference rate is *A_0_* = 2×10^9^ ATP/s-neuron, the approximate empirical value for the resting awake state estimated in Section 2.3. A value of 0.25 *A_0_* is assumed for nonsignaling functions associated with one neuron, based on the empirical results described in Section 2.3. The energy costs per neuron for the two populations are:

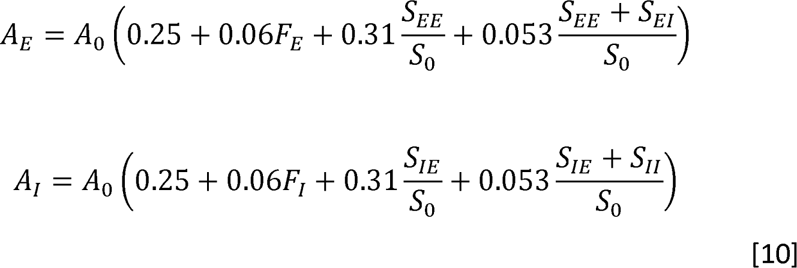

In these expressions the four terms respectively describe nonsignaling costs, the cost of spiking, the cost of excitatory synaptic currents, and the cost of presynaptic activity (which applies to both excitatory and inhibitory synapses). Taking into account the different population numbers of the two populations, the average energy use (ATP/s-neuron) of the combined populations is:

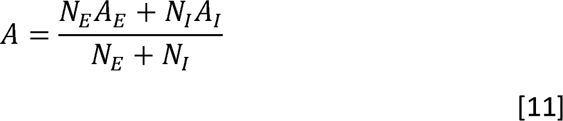

## 5. Results

### 5.1 Basic dynamics and energy costs of the WC-model

#### Model behavior illustrated with a simple numerical experiment

To compare the energy costs of model dynamics with the empirical estimate *A_0_* of ATP/s per neuron from **Figure 1**, we consider a basic experiment in which an excitatory external input *P_E_=P_E0_*is applied to the *E*-population, while the other external inputs in Eq [7] are zero. The excitatory input starts at time *t*=0 and is applied for 1s, and the resulting neural population dynamics are shown for *t* up to 1.5 s. The energy costs are estimated as the average for the period between *t*=0.5 and *t*=1.0 s, the second half of the stimulus block, and scaled to the empirical estimate *A_0_* of ATP/s per neuron from **Figure 1**. The example in **Figure 4A** illustrates the limiting case when there is no recurrent excitation or inhibition, showing a low level of evoked activity in the E-population and no activity in the I-population. The examples in **Figure 4B-D** illustrate three different types of dynamical behavior that arise from nonlinear recurrent excitation: a constant steady state, a limit cycle, and sustained activity after the stimulus has ended. For each of these examples the strength of the excitatory external input was the same (P _E0_=0.32), and the time constants for the *E*- and *I*-populations were the same ( *τ*_E_=0.01 s and *τ*_I_=0.007s). In each example, the weight *W_4_* was set to zero. The three examples of nonlinear dynamic behavior are discussed individually in the following sections.

**Figure 4.**
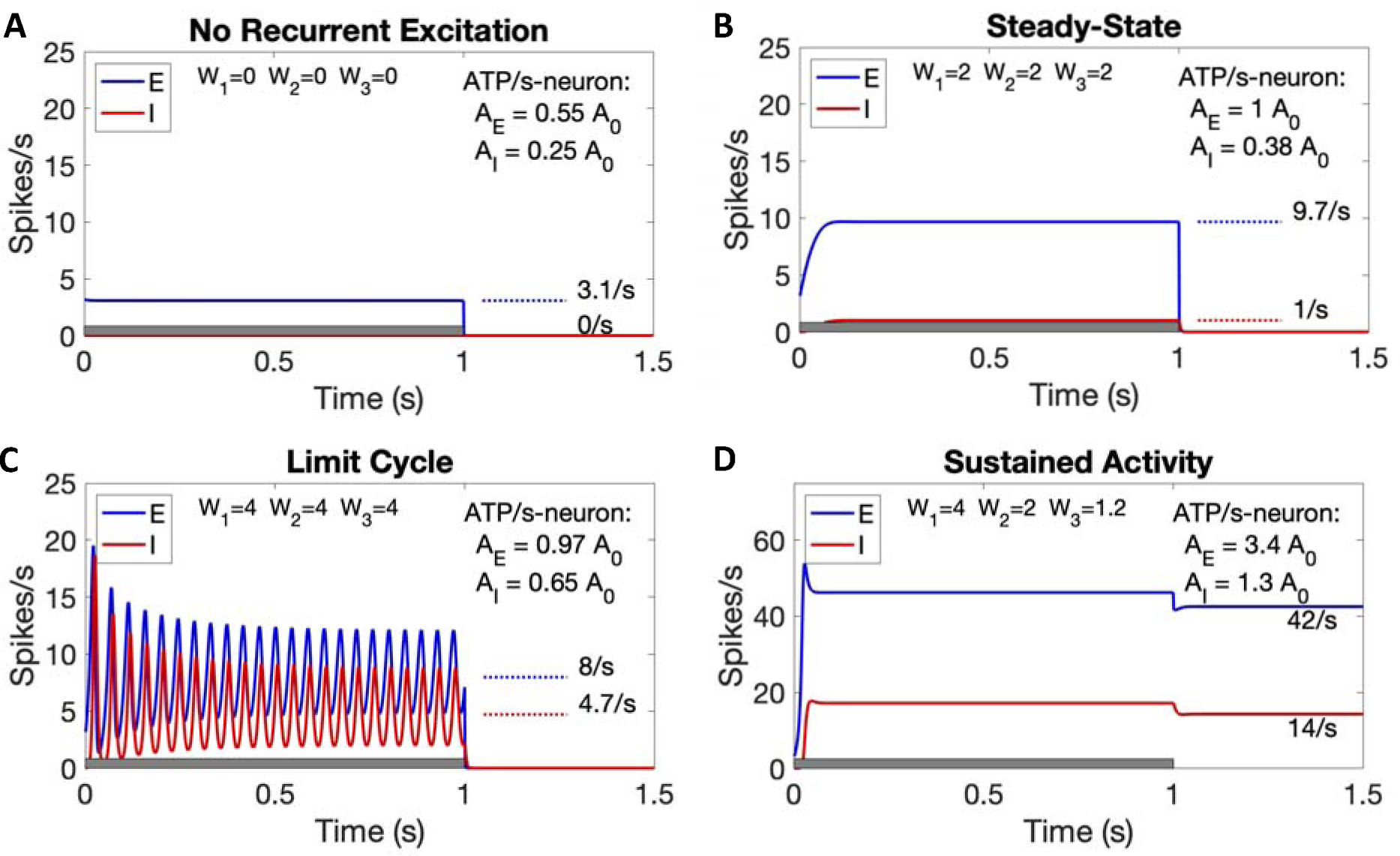
Model dynamics and rate of energy use. Results for a simple experiment in which the external excitatory input to the E-population was maintained at a constant value *P_E_*=0.32 for one second (indicated by the gray bar on the time axis). The panels differ only in the weights (W) of the internal connections. In each panel the firing rates of the *E*- and *I*-populations are shown as functions of time. For panels **A-C**, average values over the second half of the stimulus were calculated for the average firing rates of the *E*- and *I*-populations (indicated by a dotted line with the average value), and for the average energy use in ATP/s per neuron (*A_E_* and *A_I_* for the *E*- and *I*-populations). **A)** No recurrent excitatory or inhibitory activity, with all *W*’s set to zero. **B**) Steady-state behavior for balanced recurrent activity with *W_1_*=*W_2_*=*W_3_*=2 and *W_4_*=0. **C**) Limit cycle behavior when the weights are increased but remain balanced, with W_1_=W_2_=W_3_=4 and W_4_=0. **D**) Sustained activity after the stimulus ends with high recurrent excitation but reduced recurrent inhibition, with W_1_=4, W_2_=2, W_3_=1.2 and W_4_=0. Average energy use is shown for the sustained period after the stimulus. Note that the vertical scale is 3 times larger on this plot.

#### Steady-state behavior

For low values of the internal weights, the firing rate of the *E*-population reaches a steady state with constant values of *E* and *I*, with those values increasing with increasing values of the *W*’s. When the weights are all equal to zero, eliminating the recurrent excitation, the firing rate *F_E_* is about 3 spikes/s (**Figure 4A**). When the weights are increased to W _1_=W_2_=W_3_=2, *F_E_* grows due to the recurrent excitation to about 10 spikes/s (**Figure 4B**). The energy cost per neuron also increases as the weights increase, and is about equal to the reference value A _0_ for this example.

#### Limit cycle behavior

As the weights are increased further, again with W _1_=W_2_=W_3_, a new type of behavior develops. The steady-state solution becomes unstable, leading to a limit cycle in which both the *E*- and *I*-populations oscillate, creating oscillating average population firing rates (**Figure 4C**). Compared with the steady-state example for the same excitatory stimulus, the average firing rate *F_E_* is lower during limit cycle behavior but the peak firing rate of the oscillation is higher. An interesting feature of the limit cycle behavior is that the population oscillation frequency is much larger than the average firing rate *F_E_* (about 23.6 Hz for the limit cycle compared to 8 spikes/s in the example of **Figure 4C**). In these examples, both steady-state behavior and limit cycle behavior are associated with an energy use rate (ATP/s-neuron) similar to the empirical value A_0_.

#### Self-sustained high firing rate behavior

If *W_1_* is kept the same with a value of 4, but the role of recurrent inhibition is reduced ( *W_2_*=2, *W_3_*=1.2), creating recurrent excitation less constrained by inhibitory activity, the behavior is quite different (**Figure 4D**; note that the vertical scale is tripled compared to the other panels). For these weights the neural activity becomes self-sustaining, continuing after the stimulus has ended with a high firing rate ( *F_E_*=42 spikes/s) with a corresponding high energy use (*A_E_*=3.4*A_0_*). This is a point attractor, in which the state is sustained by the recurrent activity at a firing rate that is primarily determined by the recurrent activity itself rather than the input excitation. In summary, high recurrent excitation with reduced inhibition from the *I*-population can lead to sustained activity at a high firing rate after the stimulus has ended, although the energy cost per neuron is much higher than our empirical estimate.

#### Dependence of population activity on the external input

The models in **Figure 4** focused on the effects of varying the internal weights for a fixed excitatory external driver *P_E_* to the *E*-population. To test the effects of the strength of the input, the same numerical experiment with a one second stimulus was repeated for different values of *P_E_*. For each value, the average firing rates (*F_E_*and *F_I_*) and the average rate of energy use per neuron ( *A_E_*and *A_I_*) of the *E*- and *I*-populations were calculated by averaging over the second half of the stimulus. **Figure 5** shows how the firing rates (left column) and energy costs (right column) vary as P_E_ increases for the examples corresponding to the steady-state model (**Figure 4B**) and the limit cycle model (**Figure 4C**). For *W_1_=W_2_=W_3_*=2 (**Figure 5A-B**), the dynamics show steady-state behavior, with activity increasing as *P_E_*increases. For *W_1_=W_2_=W_3_*=4 (**Figure 5C-D**) the dynamics show limit cycle behavior, beginning around *P_E_*=0.3, with the range of the oscillation shown as dotted lines on the firing rate plot. Compared to the steady-state model, the limit cycle model shows a slower increase in the *average E*-firing rate with increasing excitatory input but a faster increase of the *maximum* firing rate of the limit cycle. The corresponding energy costs are plotted in the right column as a function of the excitatory input. In **Figure 5E-F** the firing rates of the *E*- and *I*-populations are plotted as a function of the average total energy use rate (Eq [11]). Note that the energy cost associated with zero firing rate is the assumed non-spiking cost of 0.25A _0_. The dotted line in each plot is a reference line corresponding to a fixed energy cost per spike of 2×10^8^ ATP, which approximately matches the *E*-firing rate curve for the limit cycle model. The steady-state model shows a steeper increase in firing rate as the energy use increases, consistent with a lower total energy cost/spike.

**Figure 5.**
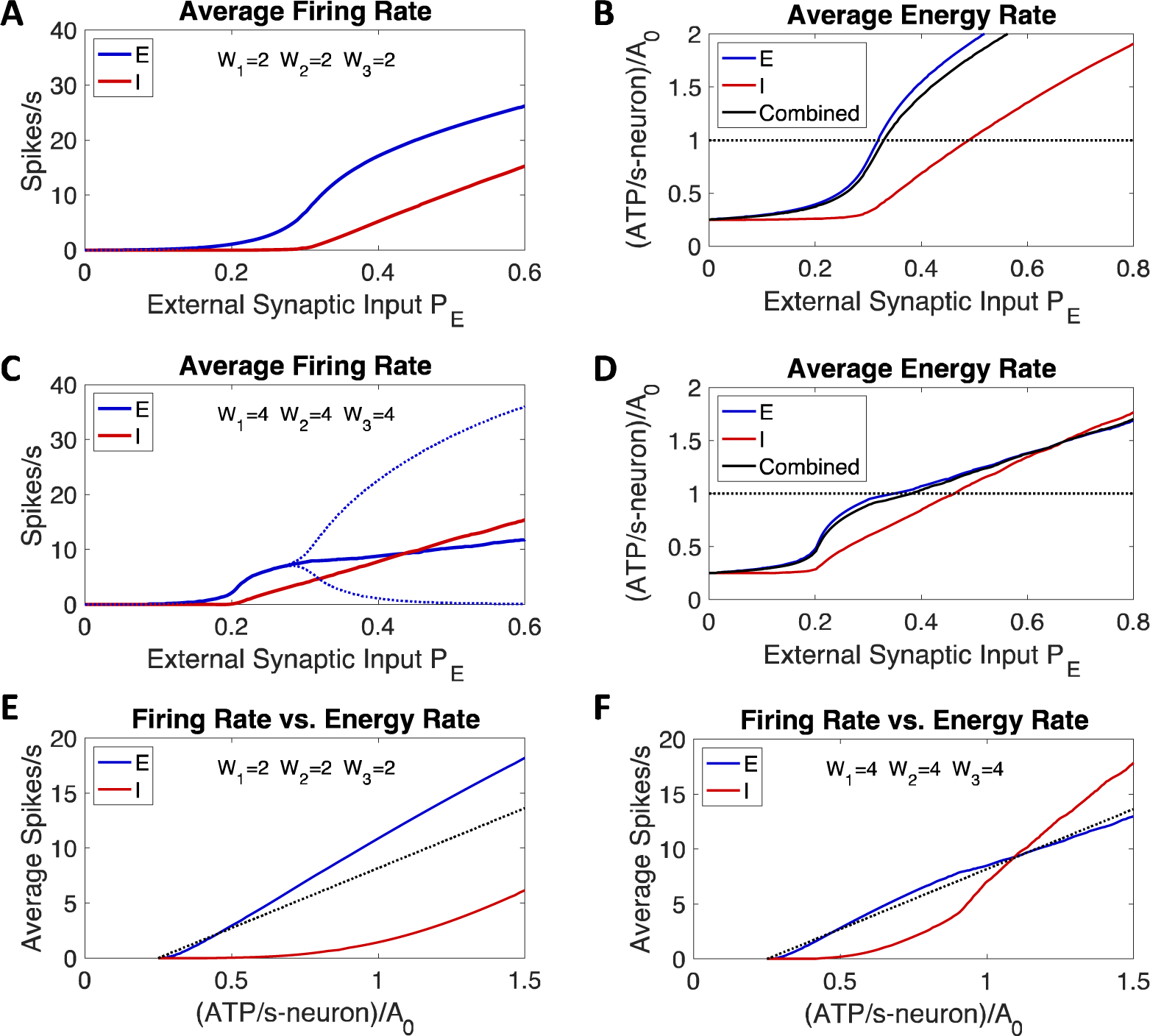
Population behavior as a function of external excitatory input. Model dynamics for steady-state and limit cycle behavior as the excitatory input to the *E*-population (*P_E_*) increases, calculated for two sets of weights. **A**) Steady-state behavior, with *W_1_=W_2_=W_3_*=2. **B**) Corresponding rate of energy use for the steady-state behavior, in ATP/s-neuron, normalized to *A_0_*, the reference empirical rate. The combined average uses Eq [11]. **C)** Limit cycle behavior, with *W_1_=W_2_=W_3_*=4. The dotted lines show the minimum and maximum values of the oscillations when a limit cycle occurs (limit cycle excursions are shown only for the*E*-population to avoid visual clutter in the figure). **D**) Corresponding rate of energy use for the limit cycle behavior. **E**) The curves plotted in **A** and **B** for the steady-state behavior are replotted to show the relation of firing rates and energy use rate. As a reference, the dotted line shows the expected result for a fixed cost per spike of 2×10^8^ ATP per spike. **F**) Similar plot from **C** and **D** for the limit cycle behavior.

#### Energetics and the frequency of the limit cycle

The oscillation frequency of the limit cycle in the example of **Figure 4C**, about 23.6 Hz, is in the range of frequencies measured empirically in EEG studies [35]. This frequency is much greater than the mean firing rate of the *E*-population, *F_E_* about 8 spikes/s, indicating that any particular neuron contributes a spike to the population average only about every third cycle. This phenomenon was highlighted by Wallace and colleagues [27] in the context of the WC-model. On the other hand, the limit cycle frequency is much less than the maximum firing rate of a single model neuron, 1/ *τ*_E_ = 100 spikes/s, or the maximum steady-state firing rate for the population as a whole, 50 spikes/s. As discussed in the **Appendix**, the limit cycle frequency depends on multiple aspects of the model, including the internal weights, the strength of the excitatory driver to the *E*-population, the time constants *τ*_E_ and *τ*_I_, and even the shape of the firing rate function *h(S)*. In terms of energetics, there is not a strong effect of the limit cycle itself; as illustrated in the examples in **Figure 4**, the energetics are primarily determined by the average firing rate, whether or not a limit cycle is produced. A general trend, though, is that as the excitatory input increases, the average firing rate increases and the limit cycle frequency also increases (see **Appendix**).

### 5.2 Dynamics of a network of interacting WC-units

#### A network of WC-units

Our basic assumption is that brain function, in all its complex forms, starts with the ability to control how neural activity spreads through a neural network, activating different neural circuits under different conditions. As a first step to illustrate how neural population activity can be controlled, and to explore the associated energy costs, we considered a simple network of interconnected WC-units. That is, rather than treating the nodes of a network as individual neurons, we now treat each node as two interacting populations of *E*- and *I*-neurons. In the diagrams to follow, a connection of the form *E_1_* to *E_2_* is indicated by a solid arrow, and a connection of the form *E_1_* to *I_2_* is indicated by a dashed arrow. We first consider one WC-unit driving another, which illustrates a classic effect in nonlinear systems: period doubling. In **Figures 6-7** we then consider a simple network of WC-nodes that functions as an ‘on-switch’, turning on to a constant persistent activity level when the input passes a threshold. The primary outcome variables of the dynamics calculations are the average firing rates *F_E_* and *F_I_*for each node, and the associated average rate of energy use *A* for each node calculated as the combined population value (Eq [11]).

**Figure 6.**
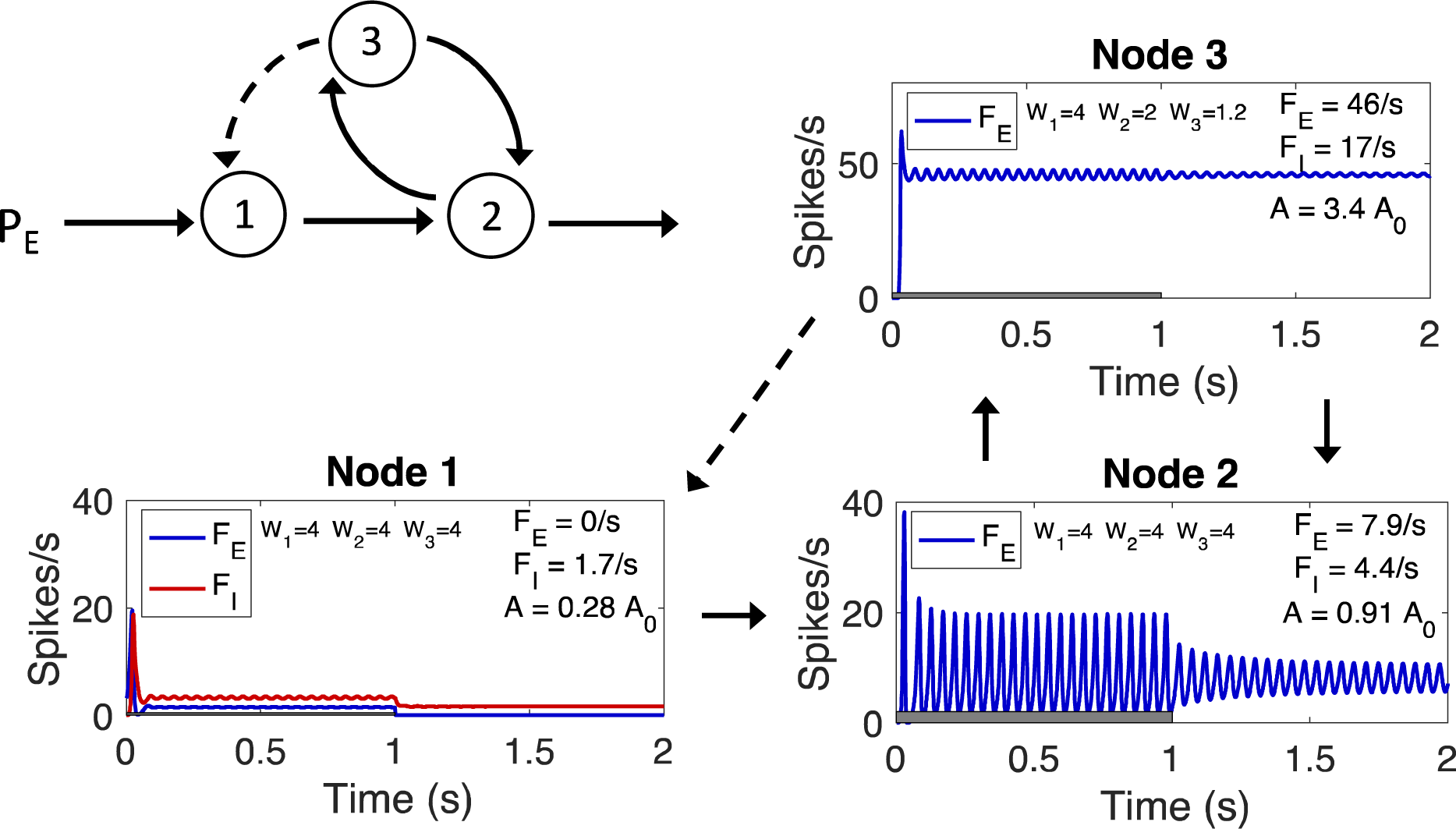
A neural ‘on-switch’. Three WC-nodes connected to function as a simple circuit element: an on-switch that starts a sustained activity in node 2, the output node that connects to the rest of a larger neural network. A schematic diagram of the connections is in the upper left, with node 1 exciting node 2, node 2 exciting node 3, and node 3 reciprocally exciting node 2 but inhibiting node 1 by exciting the I-population. The other panels show an example of the functioning of the on-switch, plotting the firing rate of the *E*-population *F_E_* of each node during the 1-s stimulus (gray bar) and after. For node 1, the I-population firing rate *F_I_* also is shown to indicate the ongoing activity of the *I*-population. Node 2 is the limit cycle example illustrated in **Figure 4C**. Node 3 is the high-firing rate fixed-point example of **Figure 4D**. Average firing rates and energy use rates are indicated for the period after the end of the stimulus, during the sustained activity.

**Figure 7.**
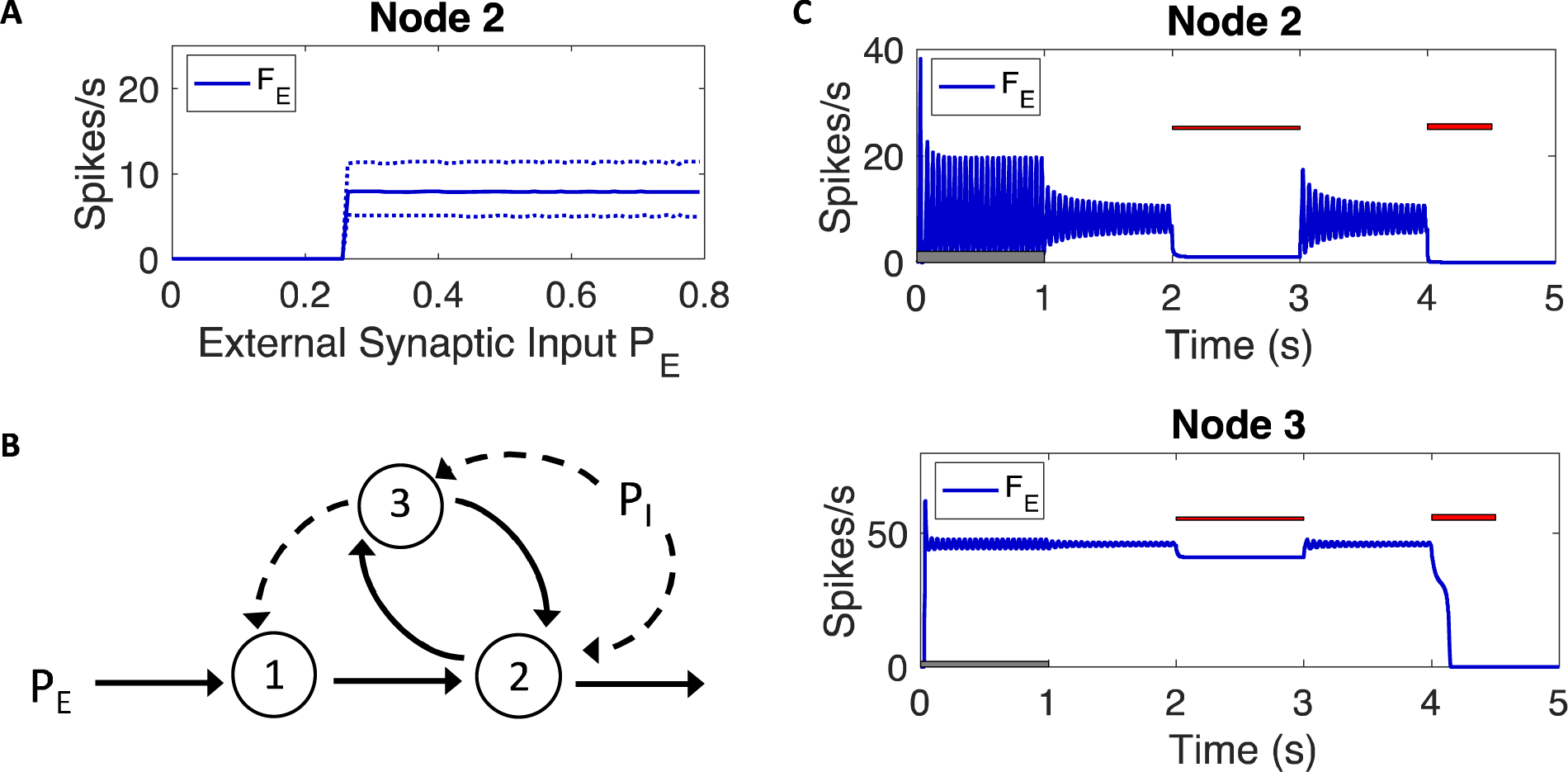
Controlling the on-switch. For the on-switch in **Figure 7**, the sustained activity of node 2, the output to the rest of the neural network, can be controlled in several ways. **A**) *Onset of sustained activity:* As the excitatory input *P_E_* to node 1 is increased the sustained activity in node 2 turns on sharply to a fixed value when a threshold is passed. The dotted lines show the maximum and minimum of the limit cycle in this example. **B**) *Inhibitory modulation*: Additional external inhibitory input *P_I_* to the *E*-populations of nodes 2 and 3 can temporarily suppress the activity in node 2 or shut the switch down entirely. **C**) *Suppression and termination*. These effects are illustrated with a more extended experiment, including initial excitatory input to node 1 to initiate the on-switch, followed by two periods of external inhibitory input to the *E*-populations of nodes 2 and 3 (red bars). Firing rates as a function of time for node 2 (top) and node 3 (bottom) are shown. During the first period of external inhibition ( *P_I_*=0.1) the output of node 2 is suppressed without killing the activity in node 3, and when the inhibition ends the sustained activity of node 2 quickly returns. During the second period of external inhibition, with stronger input ( *P_I_*=0.18), the sustained activity of node 3 is terminated and the on-switch is turned off.

#### Energetics and period doubling behavior

Frequency synchronization of different neural circuits has been hypothesized to be an important component of brain function [35–37]. In the **Appendix** we illustrate an interesting effect that arises when one WC-unit drives another. If both exhibit limit cycles when driven independently, but with different limit cycle frequencies (e.g., due to a different balance of synaptic weights), the resulting behavior of the driven node depends on the relative magnitudes of those limit cycle frequencies. As illustrated in the **Appendix**, when a node with a lower frequency drives one with a higher frequency, the frequency of the driven node matches the frequency of the driving node. However, when the roles of driver and driven nodes are reversed, so that a node with higher frequency drives a node with lower frequency, the phenomenon of period doubling appears. The frequency spectrum of the driven node shows a component at the frequency of the driver plus a component at half this frequency. Period-doubling, such as this, is a common feature of nonlinear systems (see [34] for other examples in the context of the WC-model). However, as with the earlier limit cycle examples, the energetics depends primarily just on the average firing rate underlying these population oscillations.

#### Persistent behavior: an ‘on-switch’

Many functions of the brain plausibly could be created by an attractor whose activity persists after a stimulus has ended. For example, this could be a basis for working memory, or for setting a ‘context’ switch which then affects how other inputs to the network are processed. Persistent behavior can be created by a high-firing rate point attractor (as in the example in **Figure 4D**), but the associated energy costs are high. An energy efficient unit can be constructed with three WC-nodes (**Figure 6**). The connections are such that node 1 receives excitatory external input ( *P_E_*) and drives node 2, which in turn drives node 3. Node 3 then inhibits node 1 and excites node 2. The basic dynamics are that when external input to node 1 exceeds a threshold, node 3 turns on at a high firing rate, shutting down node 1 but providing a sustained driver for node 2. The rationale of this design is that node 2 is the output node that broadcasts to the rest of the neural network, while nodes 1 and 3 are purely internal, connecting only to node 2 and not to the rest of the network. Another way to look at this model is that node 3 serves as a ‘bias current’ to node 2 that makes it self-sustaining. The reason for constructing the on-switch with node 3 is that it allows us to explicitly define the bioenergetic cost of the bias current. The key assumption for making this design energy-efficient is that the internal nodes (1 and 3) can have a lower population of neurons (this assumption is considered in more detail in the **Discussion**). With this assumption, node 3 can be a high-firing rate node with sustained activity (as in **Figure 4D**), with the combined average energy cost of all three nodes only slightly higher than our empirical reference value. In the example of **Figure 6**, if 10% of the total neurons are in node 1, 80% are in node 2, and 10% are in node 3, the overall average rate of energy use for the sustained activity is 1.1 *A_0_*.

#### Controlling the on-switch

**Figure 7A** shows the sustained firing rate of node 2 (after the stimulus ends) as a function of the excitatory input *P_E_* to node 1. The key result is that the sustained activity of node 2 turns on like a sharp switch when *P_E_*is greater than a threshold value. In **Figures 6 and 7** the sustained activity is illustrated as a limit cycle, but steady-state sustained activity could be created by replacing the weights of node 2 with those of **Figure 4B**. The sustained firing behavior of node 2 will continue indefinitely if undisturbed. However, it can be suppressed or killed by applying additional inhibitory input to nodes 2 and 3 (**Figure 7B**). **Figure 7C** illustrates the effects of external inhibitory input with a more extended experiment, still with the excitatory stimulus to node 1 from *t*=0 to *t*=1s, but now with a steady inhibitory input from *t*=2 to *t*=3s, and a shorter but stronger inhibitory pulse from *t*=4 to *t*=4.5s. The excitatory stimulus is shown as a gray bar, and the inhibitory stimuli are shown as red bars. For simplicity, we assume the inhibitory pulses are equal external inhibitory inputs to the *E*-populations of node 2 and node 3 ( *P_I_* in **Figure 3**). The panels in **Figure 7C** show the excitatory firing rate *F_E_* during the more extended experiment for node 2 (top) and node 3 (bottom). The first inhibitory input period (*P_I_*=0.1) has the effect of suppressing the output of node 2 while node 3 continues in its persistent activity state. Because node 2 is the output to the rest of the neural network, this additional inhibitory activity effectively silences the output while the inhibitory activity is present. If that additional inhibitory suppression is then removed, the activity in node 2 is re-created, driven by the still active node 3 (i.e., no fresh input to node 1 is needed to resurrect the original sustained activity). When a higher pulse of inhibitory activity (*P_I_*=0.18) is applied to both nodes 2 and 3, the fixed-point behavior is killed in node 3 and the on-switch is turned off.

## Discussion

### Scope of the work

In this work we proposed a framework for connecting the metabolic energy costs of neural activity to the variables used in neural modeling. The energy costs were based on detailed cellular estimates of the ATP required for different cellular processes involved in neural activity [10, 11]. Based on experimental studies [21, 23], an estimate of the average energy use rate, in units of ATP/s-neuron, was used as a primary comparison for the estimated energy costs of the neural models. As an example of the application of these ideas to neural models, we considered the Wilson-Cowan model [12] of two interacting populations of neurons: a larger population of excitatory neurons ( *E*) and a smaller population of inhibitory neurons ( *I*). The WC-model is a useful test case because it supports a number of features characteristic of nonlinear systems [34]. Here we estimated the energy costs of three types of attractor: simple steady-state behavior, limit cycle behavior and persistent sustained activity behavior. In addition, we explored an example exhibiting period doubling (described in detail in the **Appendix**), a phenomenon in nonlinear systems often described as a route to chaos [38]. We also began to explore how an energy efficient simple network of WC units can be constructed to act as a switch controlling the spread of activity through a network. The circuit element, composed of three WC-nodes, implements an on-switch, exhibiting a sustained limit cycle that persists after the stimulus ends provided the strength of the stimulus exceeds a threshold value.

### Primary findings

Our central finding is that the rate of energy use of a model population, calculated from Eqs [15–16], roughly follows the spiking rate of the*E*-population, with the total energy cost about equal to the empirical reference value with a spiking rate *F_E_*about 8-10 spikes/s. As a result, the energy cost is similar for steady-state, limit cycle and period doubling behaviors as long as the average firing rates are similar. In contrast, a persistent attractor involves very high firing rates, and the associated energy cost is more than 3x higher than the empirical estimate.

### Modeling the energy costs

The primary assumptions of the energy costs of different aspects of neural activity are based on detailed earlier work by Attwell and colleagues [10, 11]. The current work brings in two additional features. The first is an empirical estimate of the rate of energy use (ATP/s-neuron) based on the work of Herclano-Houzel [21] and Hyder and colleagues [23]. A remarkable feature of their work is that the energy cost per neuron is relatively independent of species. Based on this result, working with energy use as ATP/s-neuron applies across species, and this is the natural variable to consider in population models as well. It is important to keep in mind though, that connecting these estimates to measured energy metabolism, which is usually defined in terms of energy use per gram of tissue, depends on the neural density of the species. The second new feature needed to connect energy costs to neural population models was a translation of the modeled firing rate to physical synaptic currents for which the estimates of [10] could be applied. For the population model considered here, the firing rate function is proportional to a sigmoidal function *h(S)*, defined such that the firing rate is half the maximum value when the synaptic activity *S* is equal to a value *S_0_*. For estimating energy costs, our central assumption is that the synaptic current *S_0_* corresponds to an empirical physical current of 3 nA as the threshold current for generating a spike [26]. The resulting energy costs scale to higher values for a higher threshold current.

### Interpretation of synaptic and spiking energy costs

A general conclusion of the earlier estimates of the energy cost of neural activity [10, 11] was that they are dominated by the costs of post-synaptic activity, rather than spiking. Our current estimates are based on this earlier work, but with the additional feature that the spiking rates are tied to the synaptic activity through the function *h(S)*, leading to our observation that the energy costs essentially follow the spiking rate. Based on the earlier studies of the high cost of synaptic activity one might have anticipated that increasing the synaptic weights would increase the overall energy costs. However, the situation is more complicated, as illustrated in **Figure 4** (compare **Figure 4B** and **C**). The source of this phenomenon lies in the way the synaptic currents and spiking rate are interconnected: the energy cost associated with synaptic activity depends on the synaptic current, which in turn depends on the product of the synaptic weights and the average spiking rate of the population, and the spiking rate depends on the synaptic current. The net effect in **Figure 4** is that increasing the weights from those in **Figure 4B** to those in **Figure 4C** lowers the average spiking rate, and so lowers the average rate of energy use.

### Population models

A limitation of population models is that they only work with population averages, and so cannot model detailed neural circuits in which the specific neuron to neuron connections are critical. For example, neural circuits in which neuronal connectivity is specified by genetics and modified by evolution are important in many basic functions of the brain, but require more detailed modeling of individual neurons. The examples of population modeling used here are further limited to exclude transient behavior, focusing on steady-state or limit cycle responses. The population models may be most appropriate for understanding higher brain functions, such as choosing an appropriate action in different conditions and environments. A potential advantage of population models is that synaptic connections between one population and another do not require precise neuron to neuron connections. That is, if each population is individually tightly connected, external input to any of the neurons of a population will spread into population average behavior. Because of this, two populations can have reciprocal connections without any particular neuron in one population having reciprocal connections with a particular neuron in the other population. This may facilitate learning in the context of increasing connections between populations acting as units of a network.

### How many neurons are needed in a population for attractors to develop?

Neural attractors, as modeled with the WC model, are a population phenomenon. How large the population needs to be for the attractor to form is a basic question that depends on several factors, including the intrinsic variability between neurons and the variability of the response of a single neuron to the same input. In addition to the number of neurons *N* in the population, a key parameter is the connectivity of the population, defined by the probability *ρ* that one neuron connects with another neuron. Wallace and co-workers [27] investigated this question by introducing a stochastic variation during the integration of the WC equations to represent the intrinsic variability of neurons, and then compared the solutions of the stochastic model with the deterministic model (the basic WC population level equations), leading to what they termed a noisy limit cycle. They also considered a population of interacting spiking neurons, finding that noisy limit cycles could be detected in a population with *N*=1000 and *ρ*=0.1, with the limit cycles becoming more pronounced for higher connectivity. In short, their conclusion was that formation of the attractor depends on reducing the variability introduced by both the variability of the neural responses and low connectivity, which leads to higher variability of the number of AP’s arriving at a neuron for a given population size.

### Physical origin of the weights *W*

There is a second aspect to the question of population size. In the context of the current paper addressing energy use costs, the development of different types of attractor depended on the absolute values of the synaptic weights *W*. These weights are lumped parameters that depend on *N*, *ρ* and the average synaptic current *S_1_* produced by one arriving action potential. The current *S_1_* could be increased in several ways, including: increasing the number of synapses reached by an arriving AP, increasing the probability of release of a vesicle with neurotransmitter, and increasing the open time of the post-synaptic ion channel. For this reason, for a fixed population size, the corresponding model weight *W* could be increased by increasing *ρ* and/or *S_1_*. Importantly, though, for the energy use calculations only the resulting synaptic currents, determined by the weights *W*, need to be considered. For that reason, it was not necessary to specify *N*, *ρ* or *S_1_*individually in the current work.

### Neural switches

We began to consider networks in which each node is a WC-unit (i.e., each node consists of interacting *E*- and *I*-populations). The goal was to consider how a few such nodes could be connected to act as a simple circuit element that could potentially be useful for neural computation, and specifically how such circuit elements could function with energy costs consistent with the empirical average from experiments. As an example, we focused on designing an on-switch, with an output node that becomes self-sustaining at a constant level of activity (either steady-state or a limit cycle) when the input crosses a threshold value. From an energetics viewpoint, the primary challenge in the context of the WC-model is that self-sustaining activity involves high firing rates and high energy cost. However, our primary concern is with the average energy cost per neuron across the nodes of the on-switch, and if the population size of a high firing rate node is small, the total average energy use per neuron can be moderate. In the example of **Figures 6-7**, three nodes are combined with this goal in mind, with a self-sustained, high firing node as a purely internal node. That is, input from the rest of the network is only to node 1, and output to the rest of the network is only through node 2. The central assumption in terms of energy use is that node 2 has a much higher population than nodes 1 and 3, so that the high energy cost of node 3 washes out in the average energy cost across nodes. As noted above, the higher value of the weights *W* that determine the development of the attractor could be created with a small population of neurons if the connectivity or the average synaptic current produced by an arriving AP is increased. Note that the output node is not self-sustaining itself, but rather is driven by the self-sustained activity of node 3. The function of the inhibition of node 1 by node 3 is to prevent a high net excitatory input to node 2 when it is driven by both node 1 and node 3: once the self-sustained activity begins in node 3 it shuts off any further input from the rest of the network. An interesting feature of the on-switch as a circuit element is that the output (from node 2) to the rest of the network can be silenced with a low level of inhibition that is not sufficient to kill the activity of node 3. Release of that inhibition then regenerates the original sustained activity of node 2. A higher level of inhibition also kills the activity in node 3, turning off the switch.

### Neural computation exploiting nonlinear dynamics

We used the term ‘neural switch’ to describe the circuit element in the previous section because it exhibits a very strong step-like behavior: a small change in the magnitude of the input near the threshold leads to a large change in the resulting activity, but for larger inputs the resulting activity stays the same (like applying pressure to an electrical switch to turn on a light: once enough pressure is applied to flip the switch, the light stays on). In this example the strongly nonlinear behavior of the system is exploited in a controlled way to make a circuit element useful for neural computations. The resulting switch-like behavior is reminiscent of electronic circuits composed of transistors. In effect, neural switches essentially can function as *if/then* branch points in the overall network, recruiting or suppressing other neural circuits required for computational processing.

### Neural oscillations

Oscillations of neural activity are often observed, and usually interpreted as a central aspect of the functioning brain, and the possible computational benefits of oscillating neural population activity is currently an active area of investigation [35–37]. Clearly one of the interesting features of the WC-model is that oscillating behavior can develop as a limit cycle, an emergent feature of nonlinear dynamics, without the neurons themselves having intrinsic oscillatory behavior. The frequency of the limit cycle depends on multiple aspects of the model (further discussed in the **Appendix**). For considering the energetics involved, a key aspect of limit cycles is that the oscillation frequency can be much larger than the average spiking rate of the population [27]. As a result, the energy costs remain moderate because the average firing rate remains moderate, despite high population oscillation frequencies.

### Learning

In this paper we did not deal with learning: how synaptic weights are modified based on experience. The modeling framework considered here can be adapted to include a mechanism of adjusting the synaptic weights, such as spike timing dependent plasticity [39, 40]. Although weight modifications could happen at the level of internal weights of a WC-unit, the most interesting application is in the connection weights between nodes in a network of WC-units. The energy costs of the connections between nodes are included in the current modeling (i.e., each node has external inputs that are included in the energy cost estimates). In this way, the current modeling covers the ‘operational costs’ of a network, the cost of ongoing neural activity for a fixed set of weights. However, the current modeling does not include the ‘infrastructure’ costs associated with synaptic remodeling (i.e., the metabolic cost associated with increasing a synaptic weight).

### Summary and conclusions

Currently, many higher-level aspects of brain function are being approached with neural population modeling [3]. Our goal here was to attach metabolic energy costs to the synaptic activities and population firing rates considered in population models. The central question was how estimates of the energy costs might constrain and guide the development of such models to be consistent with empirical measurements. Specifically, we focused on models that exploit the nonlinear behavior of recurrent excitation, including the role of neural attractors. The calculated energy costs for population models with firing rates of 8-10 spikes/s are in good agreement with the empirical estimate of energy cost (ATP/s-neuron) derived from experimental data. In addition, in considering a network in which each node is a population model, we argued that a self-sustaining neural attractor, despite an intrinsically high rate of energy use, can be incorporated within a neural circuit element to provide sustained activity once a threshold of excitatory input activity is passed.

### Appendix

#### Changing the membrane potential with synaptic currents

The membrane potential changes as charge accumulates on the surface. This is often modeled as a simple RC circuit [25]:

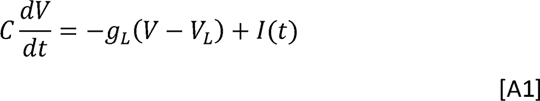

Here *C* is the capacitance of the soma membrane, and *g_L_*and *V_L_* are the conductance and reversal potential of the *leak current*. The leak current is due to a composite of ion channels open at rest that have a combined effective reversal potential equal to the resting potential *V_0_*. That is, *V_0_* = *V_L_*. If we now change variables, with *v(t)* = *V(t)* - *V_0_* as the difference of the potential at time *t* from the resting potential, and with *R* = 1/*g_L_* as the membrane input resistance, this model becomes:

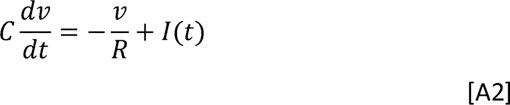

The amount of charge needed to reach the threshold potential *v* = *V_th_* (the increase over the resting potential) is determined by the capacitance *C*, and the rate of entry of charge depends on the input current *I* minus the leak current. For a constant input current *I* the time dependent voltage change is:

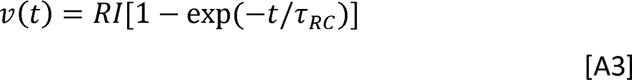

with time constant *τ*_RC_ = *RC*. For a constant current *I* that begins at time *t*=0, the maximum potential reached is *RI*. If the current *I* is too low, the threshold potential for spike generation will never be reached. The threshold current for reaching the threshold voltage is *I_th_=V_th_/R*. If *I* > *I_th_*, the time *T_th_* required to reach the threshold potential *V_th_* for a current *I* is:

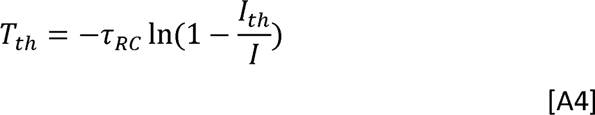

Combining Eq [3] and Eq [A4] gives an expression for the average firing rate *F*:

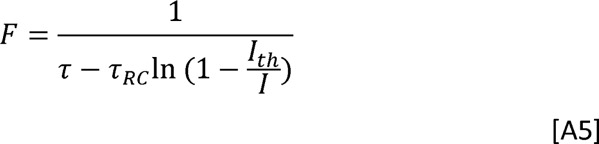

#### Assumed values for cellular properties

The firing rate (Eq [A5]) depends on three cellular parameters: the membrane resistance *R*, the membrane capacitance *C*, and the threshold voltage *V_th_*, with *τ*_RC_ = *RC*and *I_th_=V_th_/R*. For the modeling estimates we assumed a value of 0.3 nA for *I_th_* based on experimental measurements (see main text). For *V_th_*=17 mv, consistent with values reported in the literature, this implies R=57 MOhm. The capacitance *C* depends on the relevant area *A* of membrane being considered (e.g., the soma membrane area). As the area increases, *C* also increases. To remove the effect of area in specifying values, it is common to define the specific capacitance *C_m_* as the capacitance of one square centimeter of membrane (units Farad/cm^2^). The specific capacitance depends on the biophysical structure of the membrane, and has been measured to be in the range 0.7-1.0 *μ*F/cm^2^; here we assume *C_m_*= 1*μ*F/cm^2^. The relevant area of membrane *A* that must be charged to the threshold voltage to initiate firing necessarily will be uncertain due to the geometric variability of different classes of neurons. Here we take as a representative estimate *A*=1.0×10^−5^ cm^2^ (approximately the surface area of a sphere 18 *μ* in diameter). With this assumption, *C*=0.01 nF. These values for*V_th_*, *R* and *C* were used to generate the firing rate curve (blue) in **Figure 2**. To calculate the population average curve (red) in **Figure 2**, each of these parameters was randomly varied around these values with a standard deviation of 30%.

#### Variation of the limit cycle frequency

As simple tests to illustrate how the limit cycle frequency can vary, we changed one parameter at a time from the model of **Figure 4C**. Increasing the weight *W_1_* from 4 to 5 decreased the frequency from 23.6 Hz to 19 Hz. Doubling the magnitude of the excitatory external input *P_E_*, from 0.32 to 0.64, increased the frequency to 31 Hz. Decreasing the time constant *τ*_E_ from 10 ms to 7 ms increased the frequency to 28.4 Hz. The limit cycle frequency also depends on the assumed firing rate function *h(S)*. The exponent *α* in the assumed mathematical form of *h(s)* (Eq [11]) has the effect of steepening the maximum slope of the sigmoidal shape (steeper for larger *α*). Changing*α* from 3.0 to 3.5 decreased the limit cycle frequency to 18.2 Hz. Although these simple tests are not a detailed investigation of the origin of the limit cycle frequency, they indicate that the frequency depends on multiple aspects of the model. A general trend is that increasing the strength of the driving excitatory stimulus increases the limit cycle frequency, the firing rates, and the rate of energy use.

#### Frequency matching and period doubling

An interesting feature of the nonlinear dynamics emerges when one WC-unit exhibiting a limit cycle (unit 1) drives another (unit 2). If the second WC-unit has steady-state state behavior when driven with a constant input, it will oscillate at the frequency of the driver when driven by the oscillating WC-unit. When the second unit exhibits a limit cycle with a different frequency from the first when driven by a constant input, the situation is a bit more complicated. **Figure A1** illustrates two units with slightly different weights, the model of **Figure 4C**, and a model with the weight *W_1_* increased from 4 to 5. The model with *W_1_*=5 has a frequency of 17.1 Hz when driven by constant input *P_E_*=0.28, and the model with *W_1_*=4 has a limit cycle frequency of 23.6 Hz when driven by constant input*P_E_*=0.32 (top panels in **Figure A1**). When the model with *W_1_*=5 drives the model with *W_1_*=4 (that is, the lower frequency drives the higher frequency), the driven unit is matched to the frequency of the driver (**Figure A1A**). When the two units are reversed, the frequency distribution of unit 2 (now with *W_1_*=5) shows a strong component at the frequency of the driver, but in addition shows strong components at lower frequencies. These lower frequencies are consistent with period-doubling of the driver frequency, in which alternate cycles have a different amplitude. This is apparent in the lower panel of **Figure A1B**, as well as the beginning of a second period doubling. In the examples of **Figure A1**, the inputs *P_E_* to unit 1 were adjusted so the output of unit 2 was similar in mean firing rate *F_E_* and in energy cost, illustrating that the presence of period-doubling has essentially no effect on the energy cost.

**Figure A1.**
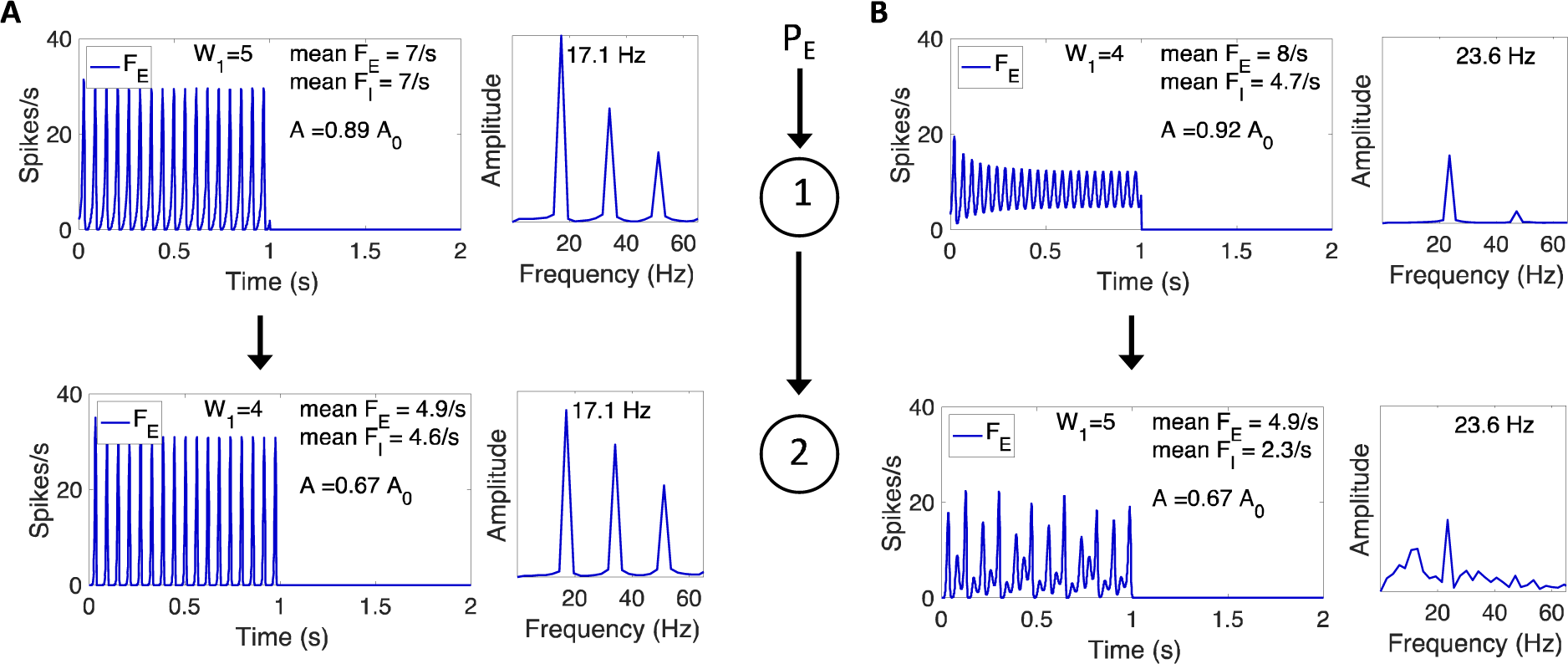
Frequency matching and period doubling. Two WC-units are arranged in a series connection with unit 1 driving unit 2 (sketched in the center). When the two units exhibit interacting limit cycle behavior, two effects can be seen: entrainment of unit 2 to oscillate with the frequency of unit 1, and a period doubling effect, with components in the unit 2 oscillation corresponding to half the frequency of unit 1. **A**) The weights for unit 1 are *W_1_*=5, *W_2_*=*W_3_*=4, exhibiting a limit cycle frequency of 17.1 Hz when driven with a constant input*P_E_*=0.28 (top panel). The weights for unit 2 (bottom panel) differ only with *W_1_*=4, and the resulting oscillations closely match unit 1. **B**) The two WC-units are reversed. Unit 1, now with *W_1_*=4, exhibits a limit cycle frequency of 23.6 Hz when driven with a constant input *P_E_*=0.32 (top panel). Unit 2, now with *W_1_*=5, exhibits a more complex spectrum, including a dominant frequency matching that of unit 1, but also additional lower frequency components consistent with period-doubling effects.

## Acknowledgements

The authors thank Thomas T. Liu and Alan Simmons for helpful discussions and thoughts on the material of this paper.

## References

1. Hawkins J, Lewis M, Klukas M, Purdy S, Ahmad S. A Framework for Intelligence and Cortical Function Based on Grid Cells in the Neocortex. Front Neural Circuits. 2018;12:121. Epub 2019/01/29. doi: 10.3389/fncir.2018.00121. PubMed PMID: 30687022; PubMed Central PMCID: PMCPMC6336927.

2. Poirazi P, Papoutsi A. Illuminating dendritic function with computational models. Nat Rev Neurosci. 2020;21(6):303–21. Epub 2020/05/13. doi: 10.1038/s41583-020-0301-7. PubMed PMID: 32393820.

3. Saxena S, Cunningham JP. Towards the neural population doctrine. Current opinion in neurobiology. 2019;55:103–11. Epub 2019/03/17. doi: 10.1016/j.conb.2019.02.002. PubMed PMID: 30877963.

4. Vyas S, Golub MD, Sussillo D, Shenoy KV. Computation Through Neural Population Dynamics. Annual review of neuroscience. 2020;43:249–75. Epub 2020/07/10. doi: 10.1146/annurev-neuro-092619-094115. PubMed PMID: 32640928; PubMed Central PMCID: PMCPMC7402639.

5. Silver D, Singh S, Precup D, Sutton RS. Reward is enough. Artificial Intelligence. 2021;299:103535.

6. Botvinick M, Wang JX, Dabney W, Miller KJ, Kurth-Nelson Z. Deep Reinforcement Learning and Its Neuroscientific Implications. Neuron. 2020;107(4):603–16. Epub 2020/07/15. doi: 10.1016/j.neuron.2020.06.014. PubMed PMID: 32663439.

7. Piloto LS, Weinstein A, Battaglia P, Botvinick M. Intuitive physics learning in a deep-learning model inspired by developmental psychology. Nat Hum Behav. 2022;6(9):1257–67. Epub 2022/07/12. doi: 10.1038/s41562-022-01394-8. PubMed PMID: 35817932; PubMed Central PMCID: PMCPMC9489531.

8. Khona M, Fiete IR. Attractor and integrator networks in the brain. Nat Rev Neurosci. 2022. Epub 2022/11/05. doi: 10.1038/s41583-022-00642-0. PubMed PMID: 36329249.

9. Buxton RB. The thermodynamics of thinking: connections between neural activity, energy metabolism and blood flow. Philos Trans R Soc Lond B Biol Sci. 2021;376(1815):20190624. Epub 2020/11/17. doi: 10.1098/rstb.2019.0624. PubMed PMID: 33190604; PubMed Central PMCID: PMCPMC7741033.

10. Attwell D, Laughlin SB. An energy budget for signaling in the grey matter of the brain. J Cereb Blood Flow Metab. 2001;21(10):1133–45. PubMed PMID: 11598490.

11. Howarth C, Gleeson P, Attwell D. Updated energy budgets for neural computation in the neocortex and cerebellum. J Cereb Blood Flow Metab. 2012;32(7):1222–32. Epub 2012/03/22. doi: 10.1038/jcbfm.2012.35. PubMed PMID: 22434069; PubMed Central PMCID: PMCPMC3390818.

12. Wilson HR, Cowan JD. Excitatory and inhibitory interactions in localized populations of model neurons. Biophysical journal. 1972;12(1):1–24. Epub 1972/01/01. doi: 10.1016/S0006-3495(72)86068-5. PubMed PMID: 4332108; PubMed Central PMCID: PMC1484078.

13. Nicholls JG, Martin AR, Wallace BG. From Neuron to Brain. 3rd ed. Sunderland, Massachusetts: Sinauer; 1992.

14. Hall CN, Klein-Flugge MC, Howarth C, Attwell D. Oxidative phosphorylation, not glycolysis, powers presynaptic and postsynaptic mechanisms underlying brain information processing. J Neurosci. 2012;32(26):8940–51. Epub 2012/06/30. doi: 10.1523/JNEUROSCI.0026-12.2012. PubMed PMID: 22745494; PubMed Central PMCID: PMCPMC3390246.

15. Buxton RB. Interpreting oxygenation-based neuroimaging signals: the importance and the challenge of understanding brain oxygen metabolism. Front Neuroenergetics. 2010;2:8. Epub 2010/07/10. doi: 10.3389/fnene.2010.00008. PubMed PMID: 20616882; PubMed Central PMCID: PMC2899519.

16. Havlicek M, Roebroeck A, Friston K, Gardumi A, Ivanov D, Uludag K. Physiologically informed dynamic causal modeling of fMRI data. Neuroimage. 2015;122:355–72. Epub 2015/08/09. doi: 10.1016/j.neuroimage.2015.07.078. PubMed PMID: 26254113.

17. Havlicek M, Ivanov D, Roebroeck A, Uludag K. Determining Excitatory and Inhibitory Neuronal Activity from Multimodal fMRI Data Using a Generative Hemodynamic Model. Frontiers in neuroscience. 2017;11:616. Epub 2017/12/19. doi: 10.3389/fnins.2017.00616. PubMed PMID: 29249925; PubMed Central PMCID: PMCPMC5715391.

18. Nicholls DG, Ferguson SJ. Bioenergetics 3: Academic Press; 2002.

19. Ames A, 3rd. CNS energy metabolism as related to function. Brain Res Brain Res Rev. 2000;34(1-2):42–68. PubMed PMID: 11086186.

20. Blaustein MP. Calcium transport and buffering in neurons. Trends Neurosci. 1988;11:438–43.

21. Herculano-Houzel S. Scaling of brain metabolism with a fixed energy budget per neuron: implications for neuronal activity, plasticity and evolution. PLoS One. 2011;6(3):e17514. Epub 2011/03/11. doi: 10.1371/journal.pone.0017514. PubMed PMID: 21390261; PubMed Central PMCID: PMCPMC3046985.

22. Herculano-Houzel S, Rothman DL. From a Demand-Based to a Supply-Limited Framework of Brain Metabolism. Front Integr Neurosci. 2022;16:818685. Epub 2022/04/19. doi: 10.3389/fnint.2022.818685. PubMed PMID: 35431822; PubMed Central PMCID: PMCPMC9012138.

23. Hyder F, Rothman DL, Bennett MR. Cortical energy demands of signaling and nonsignaling components in brain are conserved across mammalian species and activity levels. Proc Natl Acad Sci U S A. 2013;110(9):3549–54. Epub 2013/01/16. doi: 10.1073/pnas.1214912110. PubMed PMID: 23319606; PubMed Central PMCID: PMCPMC3587194.

24. Carter BC, Bean BP. Sodium entry during action potentials of mammalian neurons: incomplete inactivation and reduced metabolic efficiency in fast-spiking neurons. Neuron. 2009;64(6):898–909. Epub 2010/01/13. doi: 10.1016/j.neuron.2009.12.011. PubMed PMID: 20064395; PubMed Central PMCID: PMCPMC2810867.

25. Koch C. Biophysics of computation: Information processing in single neurons. Oxford: Oxford University Press; 1999.

26. Degenetais E, Thierry AM, Glowinski J, Gioanni Y. Electrophysiological properties of pyramidal neurons in the rat prefrontal cortex: an in vivo intracellular recording study. Cereb Cortex. 2002;12(1):1–16. Epub 2001/12/06. doi: 10.1093/cercor/12.1.1. PubMed PMID: 11734528.

27. Wallace E, Benayoun M, van Drongelen W, Cowan JD. Emergent oscillations in networks of stochastic spiking neurons. PLoS One. 2011;6(5):e14804. Epub 2011/05/17. doi: 10.1371/journal.pone.0014804. PubMed PMID: 21573105; PubMed Central PMCID: PMC3089610.

28. Destexhe A, Sejnowski TJ. The Wilson-Cowan model, 36 years later. Biological cybernetics. 2009;101(1):1–2. Epub 2009/08/08. doi: 10.1007/s00422-009-0328-3. PubMed PMID: 19662434; PubMed Central PMCID: PMC2866289.

29. Yousaf M, Kriener B, Wyller J, Einevoll GT. Generation and annihilation of localized persistent-activity states in a two-population neural-field model. Neural Netw. 2013;46:75–90. Epub 2013/05/28. doi: 10.1016/j.neunet.2013.04.012. PubMed PMID: 23708672.

30. Cowan JD, Neuman J, van Drongelen W. Wilson-Cowan Equations for Neocortical Dynamics. Journal of mathematical neuroscience. 2016;6(1):1. Epub 2016/01/06. doi: 10.1186/s13408-015-0034-5. PubMed PMID: 26728012; PubMed Central PMCID: PMCPMC4733815.

31. Wilson HR. Binocular contrast, stereopsis, and rivalry: Toward a dynamical synthesis. Vision Res. 2017;140:89–95. Epub 2017/09/09. doi: 10.1016/j.visres.2017.07.016. PubMed PMID: 28882755.

32. Chow CC, Karimipanah Y. Before and beyond the Wilson-Cowan equations. J Neurophysiol. 2020;123(5):1645–56. Epub 2020/03/19. doi: 10.1152/jn.00404.2019. PubMed PMID: 32186441; PubMed Central PMCID: PMCPMC7444921.

33. Wilson HR, Cowan JD. Evolution of the Wilson-Cowan equations. Biological cybernetics. 2021;115(6):643–53. Epub 2021/11/20. doi: 10.1007/s00422-021-00912-7. PubMed PMID: 34797411.

34. Negahbani E, Steyn-Ross DA, Steyn-Ross ML, Wilson MT, Sleigh JW. Noise-induced precursors of state transitions in the stochastic Wilson-cowan model. Journal of mathematical neuroscience. 2015;5:9. Epub 2015/04/11. doi: 10.1186/s13408-015-0021-x. PubMed PMID: 25859420; PubMed Central PMCID: PMC4388113.

35. Buzsaki G, Logothetis N, Singer W. Scaling brain size, keeping timing: evolutionary preservation of brain rhythms. Neuron. 2013;80(3):751–64. Epub 2013/11/05. doi: 10.1016/j.neuron.2013.10.002. PubMed PMID: 24183025; PubMed Central PMCID: PMCPMC4009705.

36. Uhlhaas PJ, Pipa G, Lima B, Melloni L, Neuenschwander S, Nikolic D, et al. Neural synchrony in cortical networks: history, concept and current status. Front Integr Neurosci. 2009;3:17. Epub 2009/08/12. doi: 10.3389/neuro.07.017.2009. PubMed PMID: 19668703; PubMed Central PMCID: PMCPMC2723047.

37. Cariani P, Baker JM. Time Is of the Essence: Neural Codes, Synchronies, Oscillations, Architectures. Front Comput Neurosci. 2022;16:898829. Epub 2022/07/12. doi: 10.3389/fncom.2022.898829. PubMed PMID: 35814343; PubMed Central PMCID: PMCPMC9262106.

38. Strogatz S. Nonlinear dynamics and chaos: with applications to physics, biology, chemistry and engineering: CRC Press; 2015.

39. Brzosko Z, Mierau SB, Paulsen O. Neuromodulation of Spike-Timing-Dependent Plasticity: Past, Present, and Future. Neuron. 2019;103(4):563–81. Epub 2019/08/23. doi: 10.1016/j.neuron.2019.05.041. PubMed PMID: 31437453.

40. Wong EC. Distributed Phase Oscillatory Excitation Efficiently Produces Attractors Using Spike-Timing-Dependent Plasticity. Neural Comput. 2022;34(2):415–36. Epub 2021/12/17. doi: 10.1162/neco_a_01466. PubMed PMID: 34915556.

